# SIRT6 loss causes intervertebral disc degeneration in mice by promoting senescence and SASP status

**DOI:** 10.1101/2024.09.09.612072

**Authors:** Pranay Ramteke, Bahiyah Watson, Mallory Toci, Victoria A Tran, Shira Johnston, Maria Tsingas, Ruteja A. Barve, Ramkrishna Mitra, Richard F. Loeser, John A. Collins, Makarand V. Risbud

## Abstract

Intervertebral disc degeneration is a major risk factor contributing to chronic low back and neck pain. While the etiological factors for disc degeneration vary, age is still one of the most important risk factors. Recent studies have shown the promising role of SIRT6 in mammalian aging and skeletal tissue health, however its role in the intervertebral disc health remains unexplored. We investigated the contribution of SIRT6 to disc health by studying the age-dependent spinal phenotype of mice with conditional deletion of *Sirt6* in the disc (*Acan^CreERT2^*; *Sirt6^fl/fl^*). Histological studies showed a degenerative phenotype in knockout mice compared to *Sirt6^fl/fl^* control mice at 12 months which became pronounced at 24 months. RNA-Seq analysis of NP and AF tissues, quantitative histone analysis, and *in vitro* multiomics employing RNA-seq with ATAC-seq revealed that SIRT6-loss resulted in changes in acetylation and methylation status of specific Histone 3 lysine residues, thereby affecting DNA accessibility and transcriptomic landscape. A decrease in autophagy and an increase in DNA damage were also noted in *Sirt6*-deficient cells. Further mechanistic insights revealed that loss of SIRT6 increased senescence and SASP burden in the disc characterized by increased p21, γH2AX, IL-6, and TGF-β abundance. Taken together our study highlights the contribution of SIRT6 in modulating DNA damage, autophagy and cell senescence, and its importance in maintaining disc health during aging thereby underscoring it as a potential therapeutic target to treat intervertebral disc degeneration.

## Introduction

Low back pain (LBP) is the leading cause of disability worldwide and has the highest prevalence amongst musculoskeletal conditions ^1^. The primary etiological factors for LBP are intervertebral disc degeneration and aging ^1,2^. Intervertebral disc degeneration is a progressive disease often resulting in or accompanied by spondylolisthesis and disc herniation which lead to decreased movement, pain, and disability ^2^. As the global population of ageing adults increases worldwide, it is imperative to understand the molecular basis of disc degeneration to design and develop alternate and non-surgical therapeutic approaches to confront the challenge of years lived with disability.

Sirtuins (SIRTs) are highly conserved NAD^+^-dependent histone deacetylases that function as epigenetic ON/OFF switch for genes by altering DNA accessibility and transcription^3^. There are seven homologs of mammalian SIRTs (SIRT1-7) with a discrete subcellular localizations that contribute to wider cellular processes including post-translational modifications, transcriptional regulation, energy modulation, inflammation, and cell survival^4,5^. SIRT1 is the most studied sirtuin and exerts its functions by modulating the expression and activity of key molecules such as PGC1, AMPK, and STAT^6^. Recent findings from human lifespan studies show a prominent role of nuclear Sirts including SIRT1 and SIRT6 in aging and age-associated disorders ^7,8,9^. A 2012 study by Kanfi et al. showed a significantly longer lifespan in male SIRT6 transgenic mice than wild-type mice, whereas, a recent 2021 study by Roichman and colleagues documented lifespan extension in both male and female mice, albeit with a stronger effect in males than females ^10,11^. Similar observations relating to the positive impact of SIRT6 on longevity have been reported in other species^12,13^. These studies showed that SIRT6 exerted its effect on aging through controlling activities of IGF-1 signaling and MYC pathways, both of which promote anabolic and proliferative responses by increasing cellular metabolism. SIRT6 also plays a major homeostatic role in the musculoskeletal system^14,20^. Loss of SIRT6 in osteoblast lineage cells using *Ocn-Cre* decreased osteoprotegerin expression and activated osteoclasts resulting in increased osteoclastogenesis and osteopenia ^14^. Similarly, osteoblasts and osteocyte targeted*Sirt6*^Dmp1Cre^ mice showed increased osteocytic expression of *Sost*, *Fgf23*, the senescence inducer *Pai-1,* and the senescence markers *p16* and *Il-6*, resulting in osteopenia^15^. Notably, chondrocytes derived from *Sirt6*^AcanCreERT2^ mice showed significantly hampered antioxidant defense mechanisms with decreased peroxiredoxin 1 (Prx1) levels and increased levels of an inhibitor of antioxidant activity, thioredoxin interacting protein (TXNIP)^16^. These SIRT6 loss mice presented with significantly repressed IGF-1/Akt signaling that was associated with enhanced injury-induced and age-associated osteoarthritis (OA) severity, when compared to SIRT6 intact controls ^17,16^. Similarly, recent studies of *Sirt6*^Col2a1CreERT2^ mice with SIRT6-loss in cartilage reported increased chondrocyte senescence and age-associated OA severity and showed a critical role of SIRT6 in STAT5 deacetylation which inhibited pathogenic IL-15/JAK3/STAT5 signaling^18^. Moreover, SIRT6 activation in chondrocytes prevents age-related DNA damage and suppresses senescence in an acute, traumatic disc injury model^19,20^. Collectively, these studies suggest a major role of SIRT6 in healthy aging of cartilage, bone as well as other musculoskeletal tissues. Notably, several studies also support the association of SIRT6 activity with regulation of inflammation, apoptosis, and other key mechanisms associated with aging in different tissues and model systems ^21,22^. However, despite a strong correlation of SIRT6 with aging and inflammation, the prominent etiological factors for disc degeneration, the role of SIRT6 in maintaining disc health during aging remains largely unexplored.

Herein, we investigated the role of SIRT6 in spine aging using a mouse model of conditional *Sirt6*-loss in the disc. *Sirt6* loss negatively affected disc health and the severity of degeneration increased with aging. Mechanistic studies revealed chromatin accessibility modifications primarily through modulating acetylation status of H3K9, increased DNA damage, decreased cellular autophagy and transcriptomic changes that point towards increased cell senescence and promotion of senescence associated secretory phenotype (SASP). These findings underscore a causal link between diminished SIRT6 activity and disc degeneration. Our studies for the first time highlight the critical role of SIRT6 in epigenetic regulation and maintenance of disc health *in vivo* and suggests a possible therapeutic avenue for treating age-dependent disc degeneration.

## Materials and Methods

### Mouse Studies

Animal studies were approved by the University of North Carolina and Thomas Jefferson University Institutional Animal Care and Use Committees following guidelines from the National Institutes of Health Guide for the Care and Use of Laboratory Animals. Mice were housed with an average of 4 mice per cage and had access to ad libitum water and food. Studies used male mice on C57BL/6Jx129SxFVB/NJ background since at the time of study design, prior studies demonstrated an increase in lifespan only in male mice as compared to female mice ^10^. *Sirt6^fl/fl^* mice (Jackson Labs, stock #017334) were crossed with Acan^CreERT2^ mice (Jackson Labs, stock #019148) to obtain *Sirt6^fl/fl^;Acan^CreERT2^* mice (*Sirt6*^cKO^) and littermate *Sirt6^fl/fl^* mice. At 12 weeks of age, mice of both genotypes received daily intraperitoneal injections of tamoxifen (40 mg/kg diluted to 10 mg/ml in corn oil) for 5 days. A robust deletion of *Sirt6* in Aggrecan-expressing cells in this model has been documented^14,17^.

### Histological studies

Spines were dissected and fixed in 4% paraformaldehyde (PFA) for either 6 hours or 48 hours, followed by decalcification in 20% EDTA at 4°C before embedding in OCT or paraffin for sectioning. 7μm midcoronal sections from four lumbar levels (L3-S1) of each mouse were stained with Safranin-O/Fast Green/Hematoxylin for histological assessment using a modified Thompson grading scale by at least three blind observers and with Picrosirius Red for collagen fiber characterization. Safranin-O staining was visualized using an Axio Imager 2 microscope (Carl Zeiss, Germany) using 5×/0.15 N-Achroplan or 20×/0.5 EC Plan-Neofluar objectives and Zen2TM software (Carl Zeiss).

The heterogeneity of collagen organization was evaluated using a polarizing, light microscope, Eclipse LV100 POL (Nikon, Tokyo, Japan) with a 10x/ 0.25 Pol/WD 7.0 objective and DS-Fi2 camera and images analyzed in the NIS Elements AR 4.50.00 software (Nikon, Tokyo, Japan). Under polarized light, stained collagen bundles appear either green, yellow, or red and correlate to the fiber thickness. Color threshold levels were maintained constant between all analyzed images.

### Immunohistological analyses

Deparaffinized sections following antigen retrieval or frozen sections were blocked in 5% normal serum in PBS-T, and incubated with antibodies against H3K9Ac (1:50,, Sigma, 06-942), collagen I (1:100, Abcam ab34710), collagen X (1:500, Abcam ab58632), chondroitin sulfate (1:300, Abcam ab11570); p21 (1:200, Novus NB100-1941), p-H2AX (1:50, Cell Signaling 9718), IL-6 (1:50, Novus NB600-1131), TGF-β (Abcam; ab92486) F-CHP (1:100, 3-Helix). For mouse antibodies, a MOM kit (Vector laboratories, BMK-2202) was used for blocking and primary antibody incubation. Tissue sections were washed and incubated with species-appropriate Alexa Fluor-594 conjugated secondary antibodies (Jackson ImmunoResearch,1:700). TUNEL staining was performed using the *In situ* cell death detection kit (Roche Diagnostic). Briefly, sections were deparaffinized and permeabilized using Proteinase K (20 μg/mL) and the TUNEL assay was carried out per the manufacturer’s protocol. The sections were mounted with ProLong® Gold Antifade Mountant with DAPI (Fisher Scientific, P36934), visualized with Axio Imager 2 microscope using 5×/0.15 N-Achroplan or 20×/0.5 EC Plan-Neofluar objectives, and images were captured with Axiocam MRm monochromatic camera (Carl Zeiss) and Zen2TM software (Carl Zeiss AG, Germany). Both caudal and lumbar discs were used for the analysis. Staining area and cell number quantification were performed using the ImageJ software, v1.53e, (http://rsb.info.nih.gov/ij/).

### Micro-CT analysis

Micro-CT (μCT) scanning (Bruker SkyScan 1275) was performed on fixed spines using parameters of 50 kV (voltage) and 200 μA (current) at 15 μm resolution. Images were reconstructed using the nRecon program (Version: 1.7.1.0, Bruker) and analysis was performed using CTan (version 1.17.7.2, Bruker). Transverse cross-sectional images were analyzed to evaluate trabecular and cortical bone morphology. For trabecular analysis, a region of interest (ROI) was selected by contouring the boundary between trabecular and cortical bone throughout the vertebral body. The 3D datasets were assessed for bone volume fraction (BV/ TV), trabecular thickness (Tb. Th), trabecular number (Tb. N), and trabecular separation (Tb. Sp). For cortical bone analyses, 2D assessments were computed for cortical bone volume (BV), cross-sectional thickness (Cs.Th). Disc height and vertebral length were measured at three different points equidistant from the center of the bone on the sagittal plane and used to calculate Disc height index (DHI).

### Imaging FTIR spectroscopy and spectral clustering

5 μm deparaffinized sections of decalcified lumbar disc tissues (n = 3 disc/animal, 6 animals/genotype) were used to acquire FTIR spectral imaging data using Spectrum Spotlight 400 FTIR Imaging system (Perkin Elmer, Shelton, CT), operating in the mid-IR region of 4,000 - 850 cm−1 at a spectral resolution of 8 cm^−1^ and spatial resolution of 25 μm. Spectra were collected across the mid-IR region of three consecutive sections/disc to minimize section-based variation. Using the ISys Chemical Imaging Analysis software (v. 5.0.0.14) mean second derivative absorbances in the collagen side-chain vibration (1338 cm^−1^) regions were quantified. The preprocessed spectra were used for K-means cluster analysis to define anatomical regions and tissue types within the tissue section spectral images, which represent collagen peak. Clustering images were obtained using Spectrum Image Software (LX108895).

### Tissue RNA isolation and microarray analysis

NP and AF tissues were dissected from control (*Sirt6^fl/fl^*) and *Sirt6*^cKO^ lumbar (L1-3) and caudal discs (Ca1-5). Pooled tissue from a single animal served as an individual sample. Samples were homogenized, and DNA-free, total RNA was extracted using the RNeasy® Mini kit (Qiagen). RNA with RIN > 4 was used for further microarray analysis. Fragmented biotin-labeled cDNA was synthesized using the GeneChip WT Plus kit according to the ABI protocol (Thermo Fisher). Gene chips (Mouse Clariom S) were hybridized with biotin-labeled cDNA, washed and stained with GeneChip hybridization wash and stain kit, and scanned on an Affymetrix Gene Chip Scanner 3000 7G, using the Command Console Software. Quality Control of the experiment was performed in the Expression Console Software v 1.4.1. CHP files were generated by sst-rma normalization from Affymetrix .CEL files, using the Expression Console Software. Only protein-coding genes were included in the analyses. Detection above background higher than 50% was used for Significance Analysis of Microarrays (SAM), and the p-value was set at 5%. The array data is deposited in GEO repository (GSE276439).

### NP cell isolation and treatments

Primary NP cells from adult Sprague Dawley rats (3-6 month old, Charles River), were isolated and cultured in antibiotic-supplemented DMEM and 10% FBS. To explore the role of SIRT6 *in vitro*, lentiviral particles containing Sh*Sirt6* clone #1 (RSH047819-LVRU6GP- a) and Sh*Sirt6* clone #2 (RSH047819-LVRU6GP-a) and Sh*Sirt6* clone #3 (RSH047819- LVRU6GP-a) and ShCtrl (CSHCTR001-LVRU6GP, St. Louis, MO, USA) were generated in HEK 293T cells using packaging plasmids PAX2, pRRE (#12260) and pMD2 (#12259) (Addgene, Cambridge, MA, USA) following standard protocol and stored in aliquots at - 80 °C. Primary rat NP cells were transduced with viral particles (1:1 mixture of Sh*Sirt6* clones 1 - 3 or control) with 8 mg/ mL polybrene to generate *Sirt6*-KD and *Sirt6*-Ctrl cells respectively. Medium was replaced with fresh medium containing puromycin (3 μg/ml) and after 3 days of transduction cultured in hypoxia workstation (Invivo2 400; Baker Ruskinn, UK) with a mixture of 1% O2, 5% CO2, and 94% N2 for 24 h before protein extraction to confirm the *Sirt6*-knockdown.

### Immunoblotting

*Sirt6*-Ctrl, *Sirt6*-KD NP cells were lysed and 25-40 μg of total protein was electroblotted to NC/PVDF membranes (Amersham, GE, Burlington, MA, USA). The membranes were blocked and incubated overnight at 4°C with antibodies against SIRT6 (D8D12), TXNIP (D5F3E) from Cell Signaling, and LC3 (NB100-2220, Novus). Immunolabeling was detected on the Azure 300 system using an ECL reagent (Azure biosystems, Dublin, CA) and densitometric analysis was performed using ImageJ software.

### Histone ELISA

Histone Modification Multiplex Assay kit (ab185910, Abcam) was used to quantify Histone 3 modifications including lysine acetylation and mono-di- and tri-methylation using histones isolated from Sirt6-Ctrl and Sirt6-KD cells according to the manufacturer’s instructions.

### RNA- and ATAC-Sequencing

Total DNA-free RNA was extracted from *Sirt6*-Ctrl and *Sirt6*-KD rat NP cells using RNeasy mini columns (Qiagen) (n=4 independent experiments). The extracted RNA with RIN > 7 was used for RNA sequencing. For ATAC-Sequencing, *Sirt6*-Ctrl and *Sirt6*-KD NP cells were trypsinized and collected by centrifugation at 600-800g for 5 min at 4°C. Cells were resuspended in 500 μL cryopreservation medium containing 50% serum with 10% DMSO and cryopreserved in 2 mL cryopreservation tubes. Frozen cells were shipped to Azenta for ATAC-Sequencing. The sequencing experiments were performed by Azenta using their standard protocols. The data is deposited in GEO repository (GSE276440).

### Transcriptomic data analyses using CompBio tool

Significantly up- and downregulated DEGs (FC≥ 1.5-1.75, p≤ 0.05 or FDR<0.05) were analyzed using the GTAC-CompBio Analysis Tool (PercayAI Inc., St. Louis, MO). CompBio uses an automated Biological Knowledge Generation Engine (BKGE) to extract all abstracts from PubMed that reference the input DEGs to identify relevant processes and pathways. Conditional probability analysis is used to compute the statistical enrichment score of biological concepts (processes/pathways) over those that occur by random sampling. The scores are then normalized for significance empirically over a large, randomized query group. The reported normalized enrichment scores (NEScore) represent the magnitude to which the concepts/themes are enriched above random, and an empirically derived p-value identifies the likelihood of achieving that NES by chance. The overall NEScore of ≥ 1.2 is used, resultant up- and downregulated thematic matrices are presented.

### Seahorse XF analysis

In brief, *Sirt6*-Ctrl, *Sirt6-*KD NP cells were plated in a 24-well Seahorse V7- PS test plate under hypoxia 24 hours before the experiment. On the day of experiment, cells were washed three times with 700 μl of KRPH (Krebs Ringer Phosphate HEPES) and incubated with KRPH+BSA for 1 hour at 37 °C. Seahorse XFe24 flux analyzer (Agilent Technologies) was used to determine maximum glycolytic capacity and ATP production rate using methods reported by Mookerjee et al. ^23^. Experimental design for ATP-consumption included sequential additions of 10 mM glucose, 1 μM rotenone plus 1 μM myxothiazol, 2ug/mL oligomycin. To measure glycolytic capacity, sequencial additions of 10 mM glucose, 1 μM rotenone plus 1 μM myxothiazol and 200 μM monensin plus 1 μM FCCP were performed. The normalized traces for oxygen consumption rate (OCR) and related extracellular acidification rate (ECAR) were used for calculating the experimental parameters.

### Immunofluorescence studies

*Sirt6*-Ctrl and *Sirt6*-KD NP cells were grown on poly-L lysine coated glass coverslips and fixed with ice-cold methanol for 15 minutes and blocked with 1% BSA for 1 hour. Cells were incubated with anti-LC3 antibody (NB100-2220, Novus) in a blocking buffer at 1:200 at 4°C overnight. After washing, cells were incubated with Alexa Flour 647 and mounted with ProLong Gold Antifade Mountant with DAPI. Cells were visualized using a Zeiss confocal microscope using 63x objective (CFI plan Apo Lambda 60x/1.40 oil). Staining was measured as area (pixel2 /cell) using ImageJ software (http://rsb.info.nih.gov/ij/).

### Statistics

All statistical analyses were performed using Prism7 or above (GraphPad, La Jolla). Data are represented as box and whisker plots with median, and with minimum and maximum values. Data distribution was assessed with the normality tests, and the differences between the two groups were analyzed by unpaired t-test. The differences between the three groups were analyzed by ANOVA or Kruskal–Wallis for non-normally distributed data. A chi-square (χ2) or Fischer test as appropriate was used to analyze the differences between the distribution of percentages. p ≤ 0.05 was considered a statistically significant difference.

## Results

### Sirt6^cko^ mice show accelerated disc degeneration in an age-dependent manner

The role of SIRT6 in lifespan studies has shown promising results with increased SIRT6 activity significantly extending the lifespan^10^. To study the role of SIRT6 in disc health, we characterized the age-dependent spinal phenotype of mice with *Sirt6* conditional deletion in the disc in adult mice mediated by a well-characterized *Acan^CreERT2^* allele (Fig. 1A and B). Loss of *Sirt6* increased H3K9 acetylation levels in the disc tissues confirming decreased SIRT6 levels and activity (Fig. 1C). Notably, modified Thompson grading of intervertebral discs of *Sirt6*^cKO^ mice showed significantly higher scores of degeneration in both NP and AF compartments at 12 months which became increasingly severe at 24 months when compared to their wild type littermates (Fig.1 D-F and E’-F’). The degenerative changes included NP fibrosis, focal lamellar disruptions, loss of NP-AF compartment demarcation, and clefts through NP and AF indicative of structural disruptions. Moreover, these degenerative changes in *Sirt6^cKO^* mice were more severely manifested at lower lumbar levels as compared to upper lumbar levels (Fig.1 G-G’’’). Importantly, the changes were not only evident in the lumbar spine but also in caudal discs (Suppl. Fig. 1A-B). *Sirt6*^cKO^ mice also exhibited altered disc height, vertebral height, and disc height index which is one of the indicators of disc degeneration (Fig. 1H and I- I’’)^10^. However, the vertebral trabecular bone structural parameters were only slightly affected in *Sirt6*^cKO^ mice with increased BV/TV and BMD noted only at 12 months (Suppl. Fig. 2A-E). *Sirt6*^cKO^ mice also showed some changes in the vertebral cortical bone parameters such as increased cortical porosity (Suppl. Fig. 2A, B, F). Since disc degeneration is accompanied by cell death, we performed TUNEL assay. TUNEL staining showed a slight increase in *Sirt6*^cKO^ mice, when compared to controls, without a significant decrease in total cell number suggesting that the increased cell apoptosis was not the primary driver of degeneration in this model (Fig. 1J & K-K’’).

**Figure 1:**
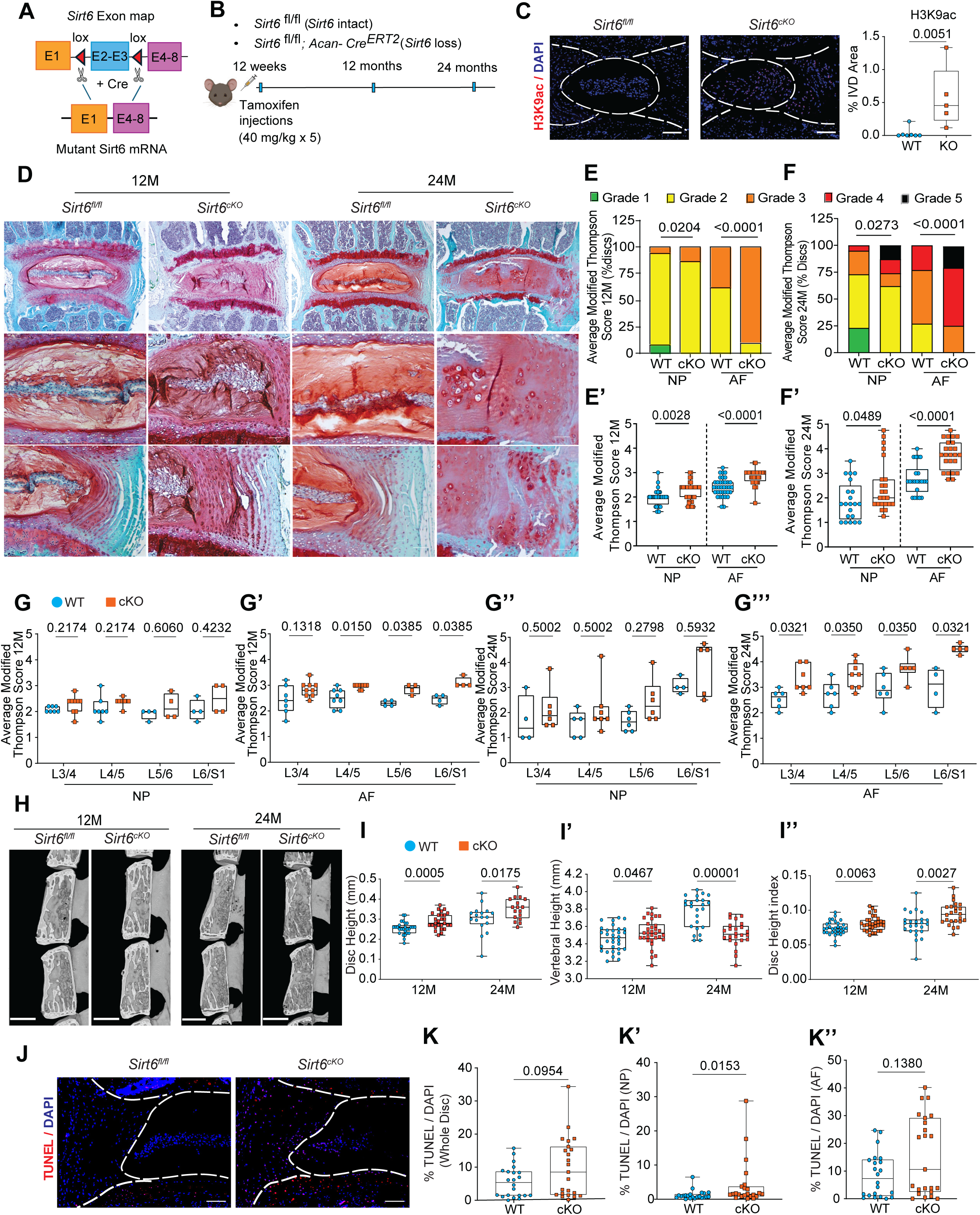
Conditional deletion of SIRT6 in intervertebral disc accelerates age-associated degeneration. **(A)** Schematic showing *Sirt6* floxed allele which following Cre-mediate recombination generates a functionally null mutant allele. (B) Experimental design showing the timeline of tamoxifen injection and analysis of control (*Sirt6*^fl/fl^) and sirt6 loss (*Sirt6^AcanCreERT2^*/*Sirt6*^cKO^) mouse cohorts. (C) Immunofluorescence staining for H3K9ac shows a robust increase in NP, AF and EP compartment of the lumbar disc confirming the deletion of SIRT6. (D) Safranin-O/Fast Green staining of *Sirt6*^fl/fl^ and *Sirt6*^cKO^ lumbar discs at 12- and 24 months. Scale bar 1A: row 1 = 200 μm; row 2 = 50 μm. (E-F) Distribution of and (E’-F’) Average Modified Thompson’s Grades of lumbar discs of *Sirt6*^fl/fl^ and *Sirt6*^cKO^ mice analyzed at 12 and 24 months. 12M: N = 4-6 mice/group, 3-4 discs/animal. 24M: N = 3-6 animals/mice, 3-4 discs/animal. (G-G’’’) Level by level Average Modified Thompson Grading scores for NP and AF compartments of lumbar discs analyzed from 12 and 24 months *Sirt6*^fl/fl^ and *Sirt6*^cKO^ mice. (H) μCT analysis showing (I-I’’) Disc height (DH), Vertebral height (VB) and Disc height index (DHI) measured at 12M and 24M. (J) TUNEL staining images and (K-K’’) TUNEL quantitation. Statistical difference between grade distributions (E-F) was tested using chi-square test, all other quantitative data was compared using unpaired t-test, *p < 0.05.

### Loss of Sirt6 affects matrix homeostasis in the disc

As Sirt6 loss accelerates disc degeneration in an age-dependent manner, we determined the matrix and cell phenotype and molecular changes in disc compartments using Picrosirius red staining, imaging-FTIR and quantitative immunohistochemistry. Picrosirius red staining showed an increased abundance of small-diameter fibers and a decrease in medium-thickness fibers in the AF of *Sirt6^cKO^* mice at 24 months indicative of dysregulated collagen turnover (Suppl. Fig. 3A, B and C). K-means clustering was used to define anatomical regions of the disc based on chemical compositions. This analysis showed similarly defined regions between *Sirt6*^cKO^ and wildtype mice at 12M. However, moderate changes in the NP compartment composition were noted at 24M, suggesting broader alterations in the disc chemical composition (Suppl. Fig. 3D-E). Notably, when the average spectra were compared, there were apparent differences in NP but not AF absorbance peaks between *Sirt6*^cKO^ mice when compared to controls at both ages, underscoring compositional differences in the NP (Fig. 3D-E). These results indicated that there were molecular changes in overall NP composition in *Sirt6*^cKO^ mice. There was also a trend of decrease in collagen-associated peak (1338 cm^−1^) in AF of *Sirt6*^cKO^ disc at 12 months which became significant at 24 months (Suppl. Fig. 3F & G). There were no differences in peaks associated with proteoglycans, between genotypes at either time point (Suppl. Fig. 3 F-G). To determine the integrity of the collagen matrix, we stained the disc sections for FCHP, a marker of denatured collagen, and COL-1. Again, *Sirt6*^cKO^ discs showed an increase in FCHP signal, along with a concurrent decrease in the abundance of healthy COL1 in the AF (Fig. 2A and B). Additionally, we observed an increase in COLX abundance in the *Sirt6*^cKO^ discs suggesting the acquisition of hypertrophic chondrocyte-like characters. There were little changes in the abundance of Aggrecan, and CS (Fig. 2A and B), suggesting that the degenerative phenotype in *Sirt6*^cKO^ was driven predominantly by altered collagen homeostasis rather than proteoglycan turnover. Overall, these results suggest that SIRT6 is critical in maintaining a healthy disc tissue matrix.

**Figure 2:**
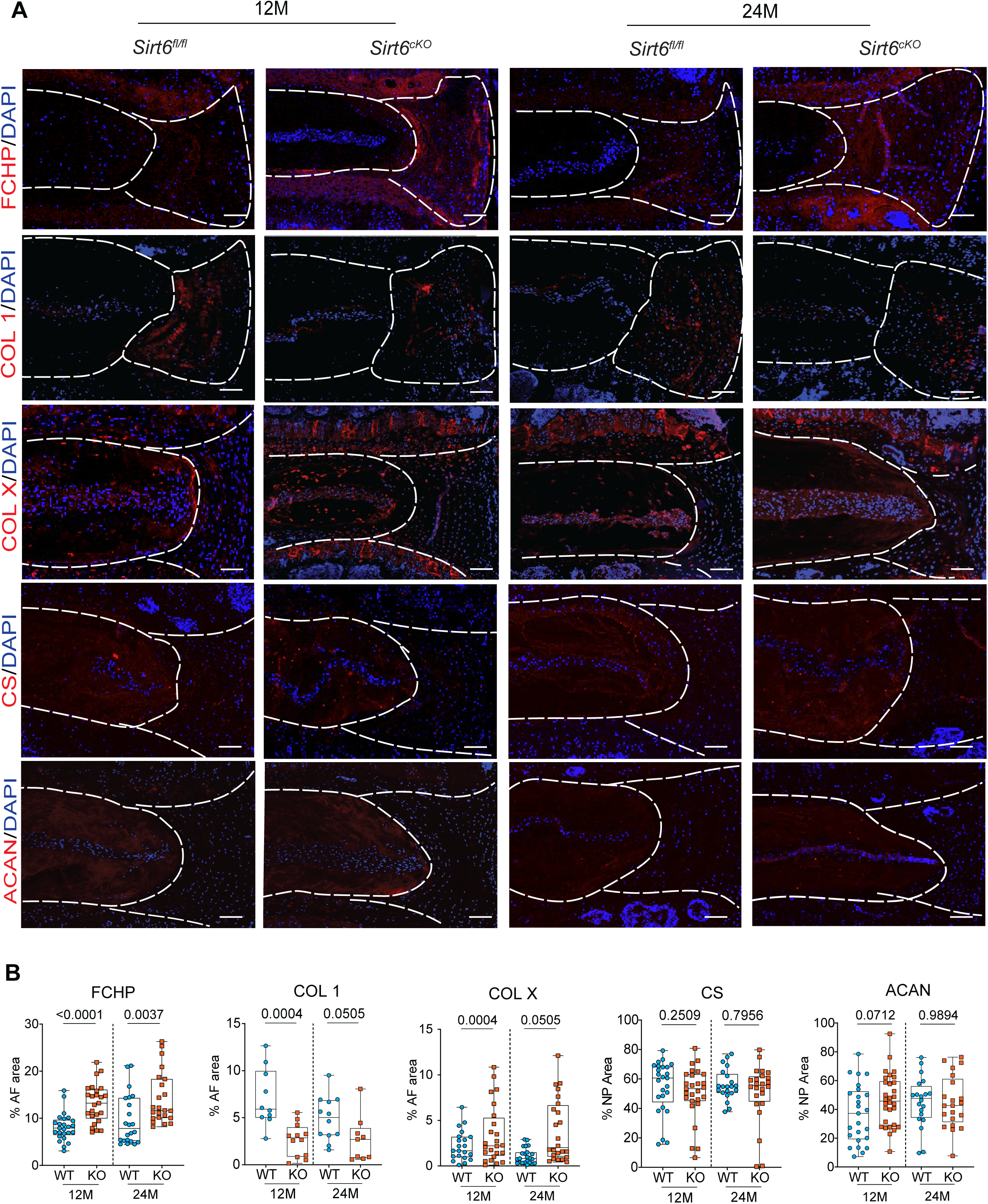
SIRT6 deletion dysregulates disc matrix homeostasis. (A) Representative immunofluorescence images of lumbar disc sections and (B) respective quantitation for FCHP, COL1, COLX, aggrecan (ACAN), chondroitin sulfate (CS) in 12M and 24M old *Sirt6*^fl/fl^ and *Sirt6*^cKO^ mice, scale bar = 50 μM. n= 4-6 mice/genotype and 3-4discs/animal. White dotted lines demarcate disc compartments. Significance was determined using unpaired t-test.

**Figure 3:**
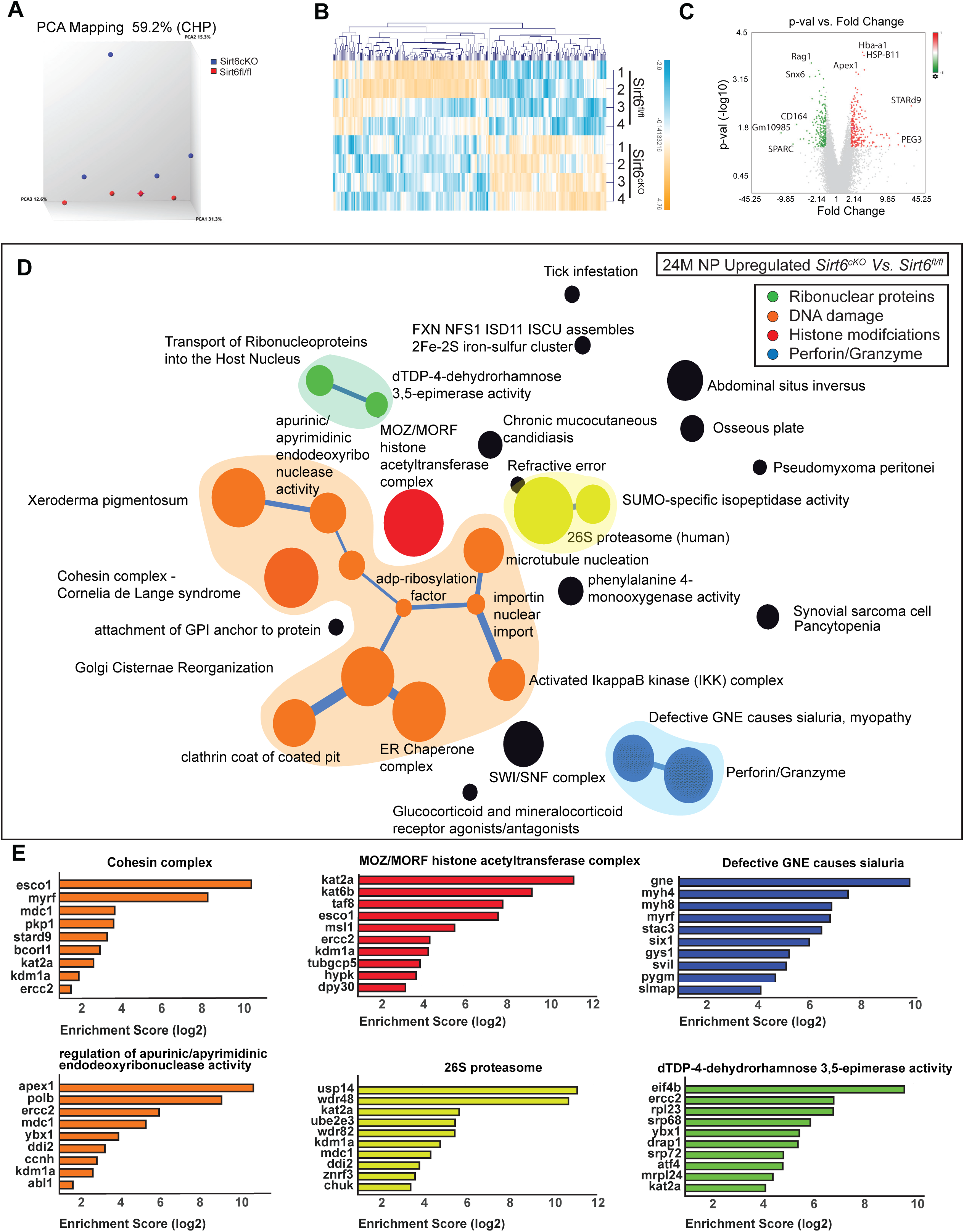
SIRT6 loss causes changes in transcriptomic landscape of NP tissues. Microarray analysis of NP tissue transcripts from *Sirt6*^fl/fl^ and *Sirt6*^cKO^ represented as (A) Three-dimensional Principal component analysis (PCA) showing discrete clustering of based on genotype (*n* = 4 mice/genotype, 5-6 pooled discs/animal) (B) Heat map and hierarchical clustering of Z-score of differentially expressed genes (DEGs) between *Sirt6^fl^*^/fl^ and *Sirt6*^cKO^ (*p* ≤ 0.05, FC≥1.75). (C) Volcano plot of DEGs in the NP showing *p*- value *versus* magnitude of change (fold change). (D) CompBIO analysis of Upregulated DEGs in NP tissue of 24M *Sirt6*^cKO^ represented in a ball and stick model. The enrichment of themes is shown by the size of the ball and connectedness is shown based on thickness of the lines between them. Themes of interest are colored, and superclusters comprised of related themes are highlighted. (E) Top thematic DEGs plotted based on CompBio entity enrichment score.

### Sirt6 loss causes major transcriptomic changes in NP and AF tissues

To delineate the molecular changes due to SIRT6 loss *in vivo*, we analyzed global transcriptomic changes in NP and AF tissues from *Sirt6*^cKO^ mice at 24 months. SIRT6 suppresses gene expression via limiting chromatin accessibility and therefore we expected to see an overall increase in gene expression in the *Sirt6*^cKO^ disc tissues. Indeed, we observed more genes were differentially upregulated in both NP and AF of *Sirt6*^cKO^ than they were downregulated as compared to their respective WT controls. In *Sirt6*^cKO^ NP, there were major upregulated thematic clusters related to i) Histone modifications ii) DNA damage iii) Ribonuclear proteins and iv) Perforins along with a smaller cluster related to proteasome (Fig. 3A-E). In the *Sirt6*^cKO^, the MOZ/MORF Histone acetyltransferase complex was upregulated (Fig. 3D & E). Another important up supercluster was the regulation of endoribonuclease activity (DNA repair), the Cohesin complex, and xeroderma pigmentosa which collectively signify an increase in DNA damage^24^. It has been widely reported that genomic instability resulting from DNA damage affects ER and Golgi ^13,14,25,26^. Accordingly, in the same cluster, we also observed connecting themes related to Golgi and ER-chaperone complex. Other major thematic clusters included changes in nuclear transport and isopeptidase activity along with IKK/NF-κB signaling which is reported to play an important role in disc degeneration^27^. In contrast to upregulated DEGs, there was no prominent thematic clustering in down regulated DEGs in the NP. Important thematic clusters upregulated in AF include DNA glycosylase, methylation, prolyl hydroxylase, protein lipidation, necroptosis, myofibrils and ABC transporter activity (Suppl. Fig. 4A-E). Again, in comparison to the extensive clustering seen in upregulated DEGs in AF, lesser clustering was observed in downregulated DEGs. Some of the enriched themes in these downregulated DEGs included BMP signaling, ubiquitin pathway, and EIFs (Suppl. Fig. 5A & B).

### SIRT6 regulates histone 3 modifications in NP cells

To delineate the mechanistic drivers of the age-dependent disc degeneration in *Sirt6*^cKO^ mice and to understand early molecular changes following *Sirt6* deletion, we performed *in vitro* loss-of-function experiments using primary rat NP cells (Fig. 4A). While SIRT6 plays an important role in various histone and non-histone modifications, its substrate binding specificity varies from tissue to tissue and its role in intervertebral disc cells is largely unknown^28^. Accordingly, we knocked down SIRT6 using lentiviral ShRNAs (Fig. 4B) and confirmed a significant decrease in SIRT6 levels and upregulation of known downstream target TXNIP (Fig.4B & C). We then probed for histone 3 modifications in knockdown NP cells. In line with our *in vivo* findings, we observed significantly increased levels of H3K9 acetylation in *Sirt6*-KD NP cells compared to cells transduced with ShCtrl (*Sirt6*-Ctrl) (Fig. 4D). Surprisingly, there were no changes in acetylation status of H3K18 and H3K56 (Fig. 4D) which are known SIRT6 targets in other cell types suggesting tissue-type specificity. Additionally, the methylation of H3K27 and H3K36 also increased in *Sirt6*-KD NP cells without changes in H3K9, H3K4, and H3K79 methylation status (Fig. 4D, Suppl. Fig. 6A).

**Figure 4:**
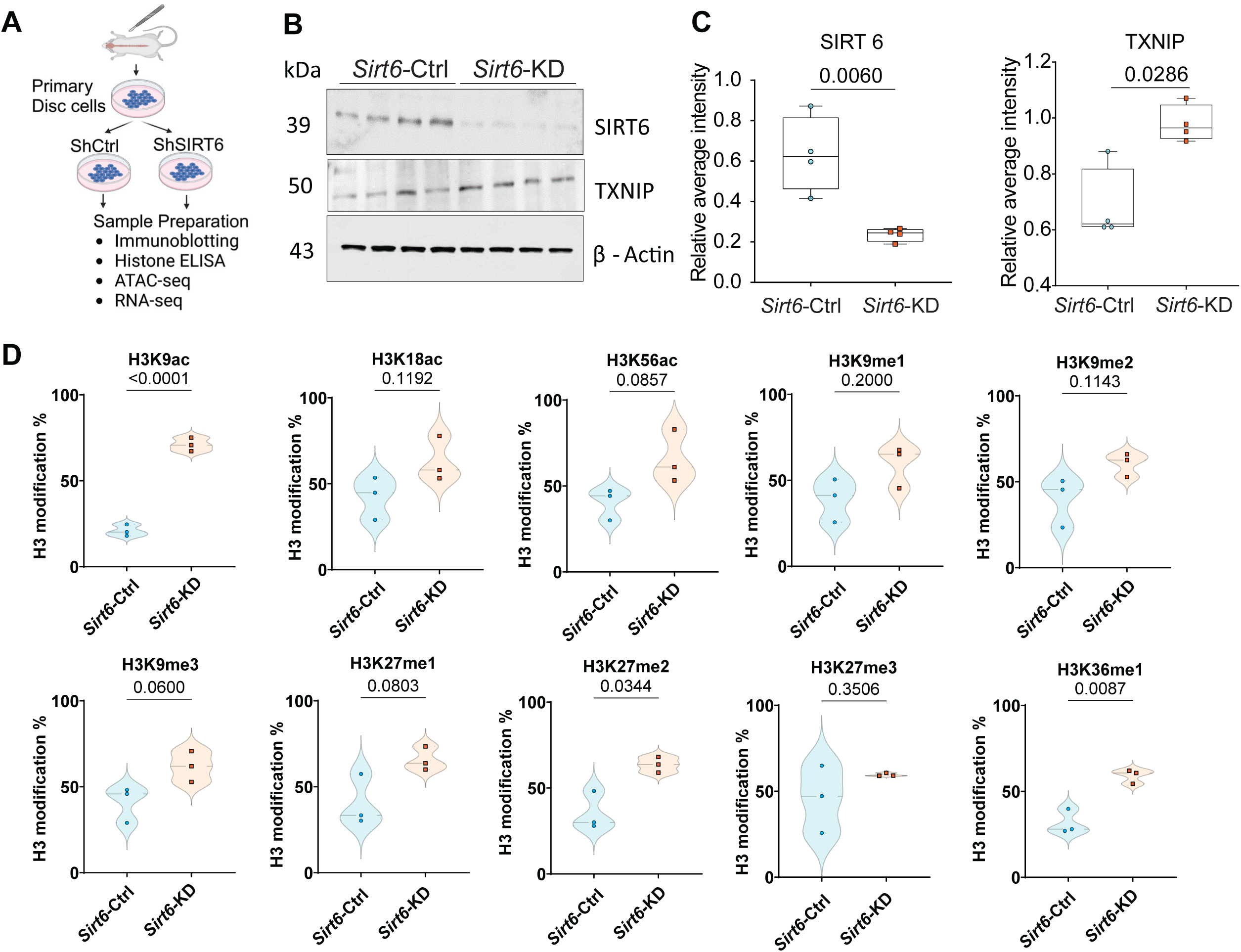
Loss of SIRT6 causes changes in histone modifications and alters chromatin accessibility. (A) Schematic of experimental design (B) Immunoblotting analysis of SIRT6 and TXNIP and (C) densitometric quantitation normalized to β-actin of *Sirt6*-Ctrl and *Sirt6*-KD NP cells. (n=4 independent cell isolations) (C) Quantitative ELISA for H3 lysine modifications from *Sirt6*-Ctrl and *Sirt6*-KD NP cells (n=3 independent cell isolations). Statistical significance was tested by unpaired t-test.

### SIRT6 knockdown in NP cells results in transcriptomic changes that align with processes affected in *Sirt6*^cKO^ mice

To determine if *Sirt6* knockdown in NP cells recapitulates transcriptomic landscape seen in 24-month-old *Sirt6*^cKO^ mice, we performed RNA-seq and ATAC-seq in *Sirt6*-KD cells. CompBio analysis of DEGs between *Sirt6*-KD Vs. *Sirt6*-Ctrl NP cells by RNA-seq revealed upregulation of five major thematic clusters i) senescence and SASP which included themes related to ECM and cytoskeletal remodeling, TGF-β and cytokine signaling, ii) BMP signaling iii) chondrosarcoma and heparan sulfate N-deacetylation iv) abnormal collagen deposition and IKK/NF-κB signaling, and v) SLIT and ROBO, mossy fibers and spinocerebellar ataxia type I (Fig. 5A-D). The downregulated clusters in *Sirt6*-KD cells included DNA damage repair pathways, sterol demethylase, reticulo spinal tract processes, cancellous bone, cartilage, and paraxial mesoderm (Suppl. Fig. 7A and B). Again, many of these processes, for example defect/decrease in DNA damage repair, have been widely correlated with degenerative and senescence phenotypes in multiple tissues^29,30^. We then determined overlapping upregulated and downregulated themes between the *Sirt6*-KD NP cells and *Sirt6*^cKO^ NP tissue using an Assertion Engine module within CompBio. A significant overlap in upregulated themes between the two datasets were noted for ECM proteins, myofibroblasts/tissue fibrosis, cell-cell, cell-matrix adhesion, actin cytoskeletal, endocytic vesicles, cartilage and arthritis, and synovitis ^31,27,32^. Within downregulated datasets, themes related to lipids/fatty acids, steroid hormones, endochondral processes, proteoglycans and paraxial mesoderm differentiation were shared (Suppl. Fig. 8A and 9A).

**Figure 5:**
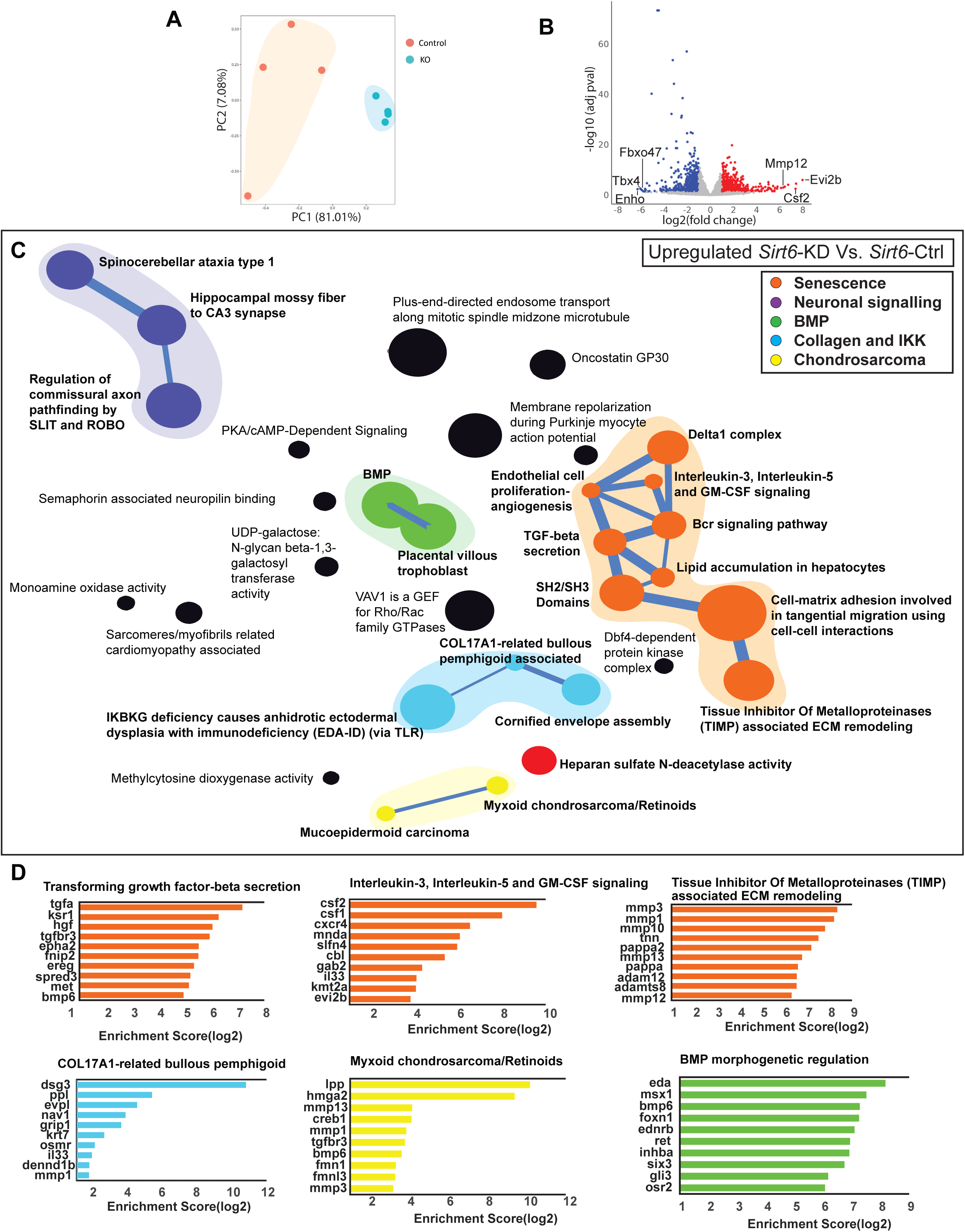
*Sirt6* knockdown in NP cells causes transcriptomic changes in senescence and ECM related pathways. RNA-Sequencing of *Sirt6*-Ctrl and *Sirt6*-KD represented as (A) Three-dimensional PCA showing discrete clustering of samples based on the genotypes (*n* = 4 independent samples/ group) (B) Volcano plot of DEGs from *Sirt6*-KD vs *Sirt6*-Ctrl. (C) CompBio analysis for upregulated DEGs (FDR<0.05, FC>1.5) represented as a ball and stick model. Themes of interest are colored, and superclusters comprised of related themes are highlighted. (D) Top thematic DEGs plotted based on CompBio entity enrichment score.

Next, we checked for common genes between ATAC- and RNA-seq datasets, these are shown in a heatmap and a quadrant plot with representative average chromatin accessibly maps for a select gene (Fig. 6A-C). We found several commonly upregulated genes including *Csf1*, *Tgfa*, *Cdk6, Bet1,* and *Itga3*, *Dock4*, *Akap6* (Fig. 6A-C). Notably, the TGF pathway is a well-known marker of senescence across various tissues^33,34^. The commonly downregulated genes in ATAC and RNA seq included *Tbx4*, *Scd*, *Sqle*, *Dcn*, *Cys1, Egflam*, *Cdh20*, *Kbtbd8*, indicating a decrease in lipid metabolism (Fig. 6A-C).

**Figure 6:**
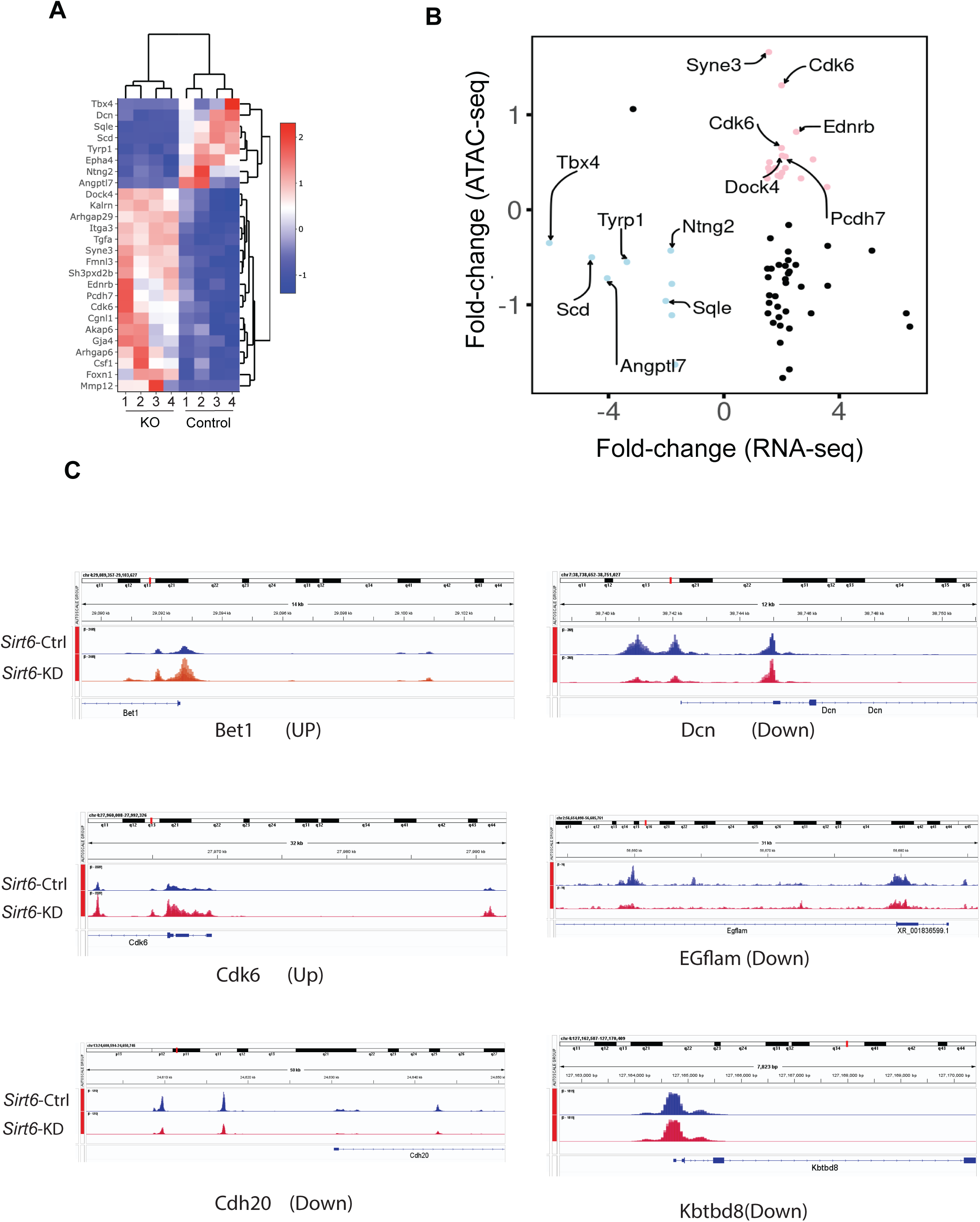
Loss of SIRT6 results in changes in chromatin accessibility. (A) Heatmap and (B) Quadrant map of commonly upregulated and downregulated DEGs (FDR<0.05, FC>1.5) between ATAC-seq and RNA-seq mapped to one of the thematic superclusters in RNA-Seq data. (C) Gene tracks showing the enriched peaks for a select group of genes from ATAC-seq experiment. (n=4 independent samples/group).

Our transcriptomic analyses hinted at downregulation in metabolic parameters of NP cells. SIRT6 has been previously shown to affect glycolysis and metabolism in mouse embryonic stem cells ^35^. Interestingly, however, measurements of 2DG uptake showed no differences suggesting a lack of altered glycolytic flux in *Sirt6*-KD NP cells (Suppl. Fig. 10A-E). We further confirmed this finding using seahorse assays to determine whether *Sirt6*-KD cells exhibit changes in glycolytic capacity and ATP production. It was evident that glycolytic capacity and ATP yields of *Sirt6*-KD cells were comparable with *Sirt6*-Ctrl cells implying a lack of effect on the glucose metabolism of NP cells (Suppl. Fig. 10A-E).

### SIRT6 loss increases senescence and SASP burden in disc

Transcriptomic data in *Sirt6*^cKO^ mice and *Sirt6*-KD cells suggested changes in cell senescence signaling pathways along with the potentially related effect on DNA repair pathways and the endocytic/autophagic pathway, which are important drivers of accelerated aging ^36^. Senescence is often accompanied by the SASP and has been linked to disc degeneration ^37^; we therefore investigated senescence signatures in *Sirt6*^cKO^ mice. Our findings showed that p21 levels were significantly increased in the discs of *Sirt6*^cKO^ mice (Fig. 7A) ^38^. Additionally, we determined levels of IL-6 and TGF-β, the primary SASP markers in the disc ^37^. Levels of both IL-6 and TGF-β, were significantly elevated in *Sirt6*^cKO^ discs (Fig. 7A) There was also an increased accumulation of lipofuscin, one of the primary hallmarks for senescence, in *Sirt6*^cKO^ discs (Fig. 7C). Since senescence is a gradual process caused by accumulation of DNA damage or alterations in autophagy, we measured primary markers for both of these processes in *Sirt6*^cKO^ mice and Sirt6-KD cells. Notably, staining for γH2AX, a marker of DNA damage was increased in the *Sirt6*^cKO^ discs (Fig. 7D). SIRT6 has been shown to modulate autophagy in various tissues^39^ which plays a key role in disc health^40^. The loss of SIRT6 decreased LC3 levels in NP cells (Fig. 7E-F & G-H) suggesting dysregulated autophagy. Together, these results suggest that increased cell senescence and SASP burden with DNA damage and dysregulation in autophagic and ER/Golgi pathways, in part, drive the degenerative phenotype seen in *Sirt6*^cKO^ mice.

**Figure 7:**
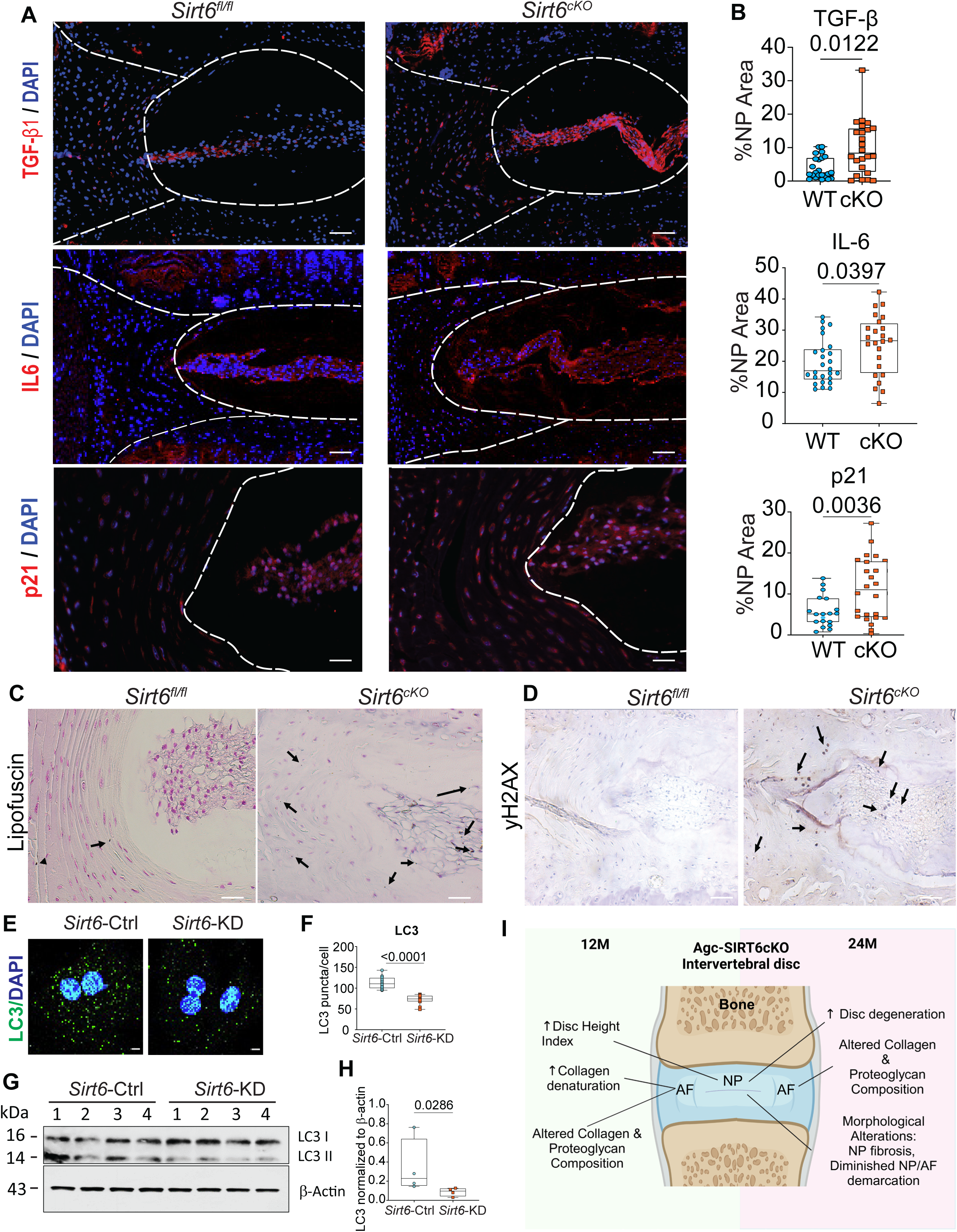
SIRT6 deletion increases DNA damage, senescence and SASP burden in *Sirt6^cKO^* discs and modulates NP cell autophagy. (A) Representative immunofluorescence images and (B) quantitative analysis of SASP markers IL-6, TGF-β, p21. n= 6 animals/genotype, 3 discs/mouse. Scale bar = 50 μM (C) Representative images of Sudan Black staining of intervertebral discs shows increased Lipofuscin accumulation in *Sirt6*^cKO^ compared to *Sirt6*^fl/fl^. (D) Representative immunostaining images of yH2AX in *Sirt6*^cKO^ and *Sirt6*^fl/fl^ intervertebral discs, C, D: n = 3 animals/genotype, 1-2 discs/mouse. (E) Immunofluorescence analysis and (F) quantitation of LC3 puncta in *Sirt6*-KD and *Sirt6*-Ctrl cells cultured in hypoxia. Significance was tested with unpaired t-test (G) Immunoblot of LC3 and (H) densitometric quantification showing LC3II/actin from *Sirt6*-KD and *Sirt6*-Ctrl cells under hypoxia. (I) Schematic showing age dependent consequences of SIRT6 loss in the spine.

## Discussion

Studies have shown a strong correlation between SIRT6 and aging, and *Sirt6* is significantly associated with increased lifespan in mice^10,11^, in centenarian humans^41^ as well as in long-lived species^42^. Despite a prominent effect on skeletal tissues^17,18^ and aging, which is one of the major risk factors for disc degeneration, the role of SIRT6 in disc health has been largely undermined^43^. Here, we show for the first time that conditional deletion of *Sirt6* in the mouse intervertebral disc significantly accelerates degeneration and promotes a severe senescent phenotype. Importantly, these changes start as early as 12 months of age thereby demonstrating a critical role of SIRT6 in disc health during aging. This study not just establishes a direct correlation between SIRT6 and intervertebral disc health but also provides the multiple pathways regulated by SIRT6, which can be explored for further studies in musculoskeletal disorders as well as in aging.

While SIRT6 affects various signaling pathways and cellular processes in different tissues, its functions can be tissue specific. Previous studies have shown that loss of SIRT6 in mouse cartilage (*Sirt6*^AcanCreERT2^) diminishes pro-anabolic IGF-1 and AKT signaling in articular chondrocytes and results in an increase in injury-induced and age-associated knee osteoarthritis^17^. Similarly, in *Sirt6^Col2CreERT2^* mice, articular chondrocytes show activation of the proinflammatory IL15/JAK3/STAT5 signaling axis which enhances OA severity^18^. However, given the unique anatomical and avascular nature of the intervertebral disc, it is plausible that in addition to the canonical mechanisms, SIRT6 may govern disc function through the modulation of pathways that are tissue specific. Our studies showed that *Sirt6* deletion in disc cells significantly increased acetylation of H3K9, along with methylation levels of H3K27 and H3K36 thereby affecting chromatin accessibility and broader gene expression changes. Importantly, we noted a substantial overlap in enriched themes between the *in vivo* (*Sirt6*^cKO^) and *in vitro* (*Sirt6*-KD) transcriptomic datasets with select upregulated themes related to ECM proteins, myofibroblasts/tissue fibrosis cell-cell and cell-matrix adhesion, actin cytoskeletal, endocytic vesicles/autophagy, FGF, vasculogenesis, cartilage/chondrogenesis, arthritis, and synovitis suggesting dysregulated ECM homeostasis, osteochondral pathways and inflammation, all processes linked to disc degeneration^31,27,32^. To this effect, we observed that *Sirt6*^cKO^ discs showed diminished Col1 abundance and increased FCHP binding suggesting altered ECM dynamics and an overall increase in collagen denaturation. As a likely compensatory response, there was an increase in thin collagen fibers suggesting stimulation of collagen turnover, and fibrosis in the NP, a known hallmark of disc degeneration^44^. Similarly, downregulated shared themes were related to lipids/fatty acids, steroid hormones, proteoglycans, and paraxial mesoderm differentiation indicating altered lipid/sterol signaling and cell differentiation. Collectively, an increase in DNA damage and a concomitant decrease in autophagy along with changes in several ECM-associated genes contributed to the degenerative phenotype observed in *Sirt6*^cKO^ mice.

Autophagy has been reported to contribute to DNA damage-induced senescence^45^. Indeed, this was supported by a decrease in LC3 levels and increased abundance of y- H2AX, p21, and accumulation of lipofuscin in *Sirt6*^cKO^ mice. Furthermore, higher levels of the known SASP markers, IL-6 and TGF-β, in *Sirt6*-deficient disc cells underscored their senescent phenotype and role of SIRT6 in senescence inhibition in disc ^46^. Our findings are in line with a report showing rescue of acute injury-induced disc degeneration by SIRT6-overexpression through modulation of autophagy and senescence ^20^. Interestingly, previous studies have shown a lack of TGFβ-regulation by SIRT6 in human fibroblasts^47^, signifying the tissue-specific role of SIRT6 in modulating this pathway. Collectively, these results establish the role of SIRT6 in counteracting intervertebral disc senescence with aging. Our results are also concurrent with the recently published information theory of aging which measures aging as a factor of epigenetic changes^48^.

Hyperacetylation of H3K9 in promoters of *Runx2*, *Osx*, *Dkk1* and *Opg* in young SIRT6 knockout mice results in low turnover osteopenia by affecting both osteoblastogenesis and bone resorption^49^. Moreover, *Sirt6* deletion has deleterious effects on articular chondrocytes and shows alterations in proliferating and hypertrophic zones of the growth plate^17,18^. Since lineage tracing studies have shown that the *Acan*^CreERT2^ allele also targets EP and growth plate cartilages^50^, it prompted us to investigate whether deletion of *Sirt6* shows any changes in vertebrae. Notably, *Sirt6*^cKO^ mice showed a small increase in bone volume, and vertebral height at 12 months, suggesting accelerated metaplasia/differentiation of hypertrophic chondrocytes into osteoblastic cells^51,52,18^.

However, these early gains in bone mass and vertebral height were followed by a decrease in vertebral height at 24 months, suggesting dysregulated growth plate dynamics and cell exhaustion with aging^53,18^. Interestingly, increased COL10 expression and acquisition of hypertrophic chondrocyte morphology by NP cells further supports the notion that SIRT6 loss promotes acceleration of the cell differentiation program in the spine.

In summary, our studies for the first time establish a causal and positive relationship between SIRT6, a nuclear histone deacetylase, and disc health *in vivo*. SIRT6-loss promotes disc degeneration by negatively regulating several key molecular and cellular processes such as ECM homeostasis and autophagy and by promoting aberrant cell differentiation, DNA-damage and senescence. Modulating SIRT6 activity using specific drugs may therefore offer an attractive, non-invasive strategy to ameliorate age- dependent disc degeneration and to preserve disc health in the aging spine.

## Acknowledgments

We would like to thank Kathryn Kelley for mouse colony maintenance and tissue collection and Dr. Andrzej Steplewski for help with FTIR spectroscopy. This study was supported by the Michael Michelson Gift Fund and NIA grants R01AG073349 (M.V.R.), R01AG044034 (R.F.L.), and R01AG078609 (J.C.). Some aspects of this research were conducted while J.C. was an Irene Diamond Fund/AFAR Postdoctoral Transition Awardee in Aging.

## Author Contributions

P.R., J.C., R.L., and M.V.R. conceptualized, conceived and designed the experiments. P.R., B.W., M.T., V.T., S.J., M.T., performed the experiments, collected, and analyzed the data. P.R, R.A.B. and R.M. performed bioinformatics analysis.

P.R. and M.V.R. interpreted the results and wrote the original draft of the manuscript. All authors reviewed and approved the final draft of the manuscript.

## DATA AVAILABILITY

RNA microarray and RNAseq data associated with this study are deposited in the GEO database with accession # GSE276439 and GSE276440. All datasets generated and analyzed during this study are included in this published article.

## ETHICS STATEMENT

All animal experiments were performed under IACUC protocols approved by the University of North Carolina at Chapel Hill and Thomas Jefferson University.

