## Supplementary figures and images for "SIRT6 loss causes intervertebral disc degeneration in mice by promoting senescence and SASP status"

### Supplementary Figure 1

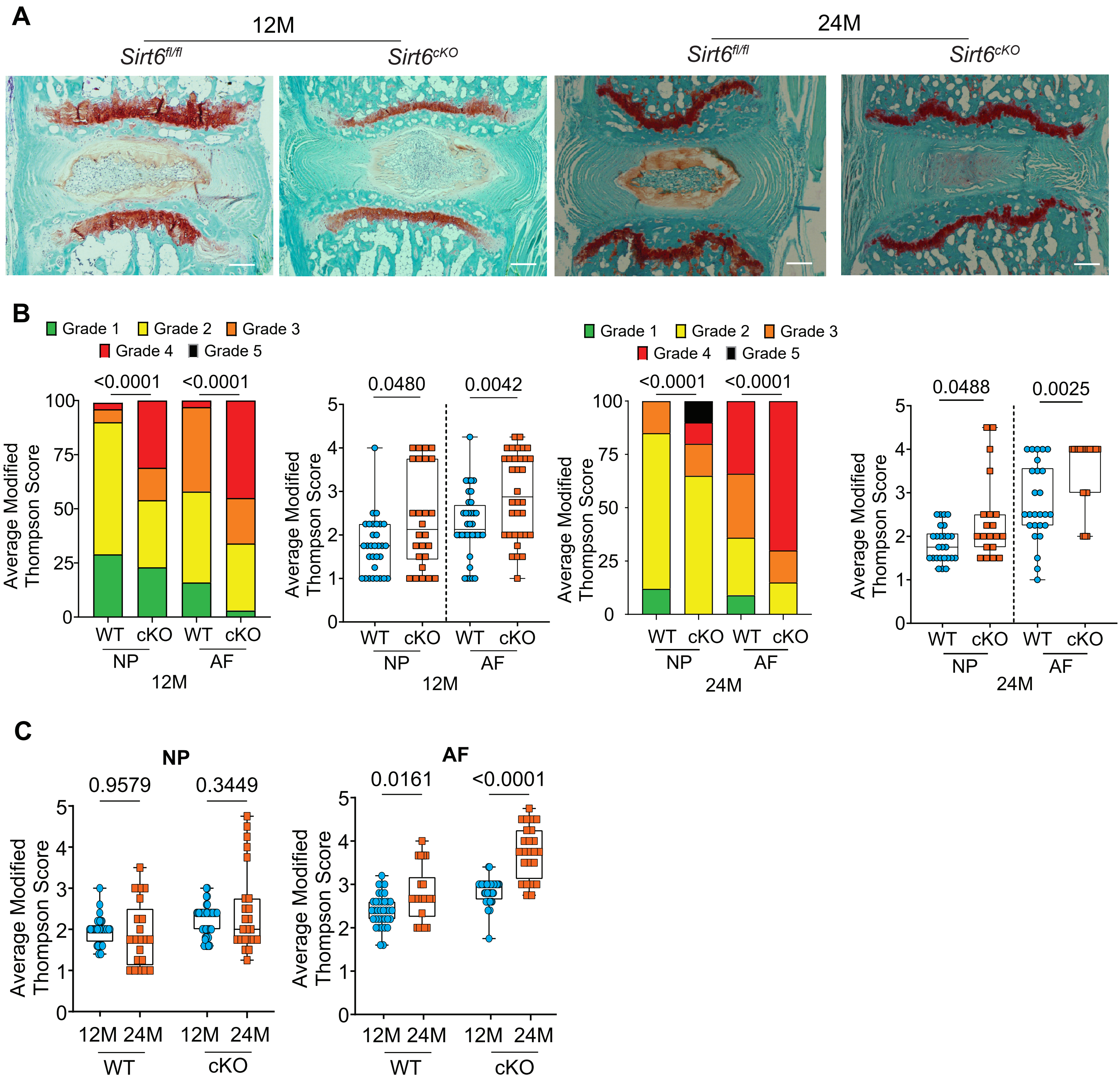

### Supplementary Figure 2

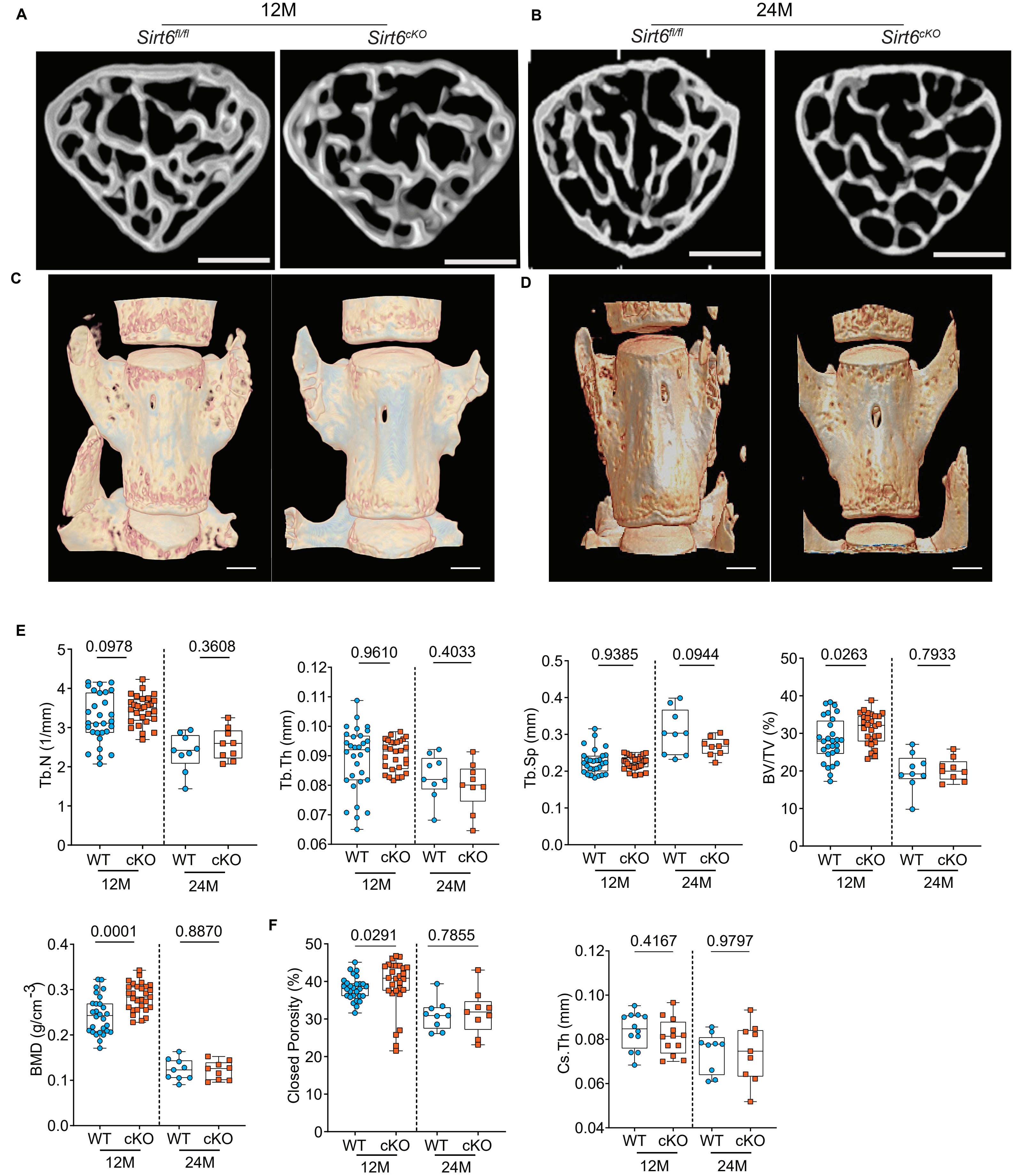

### Supplementary Figure 3

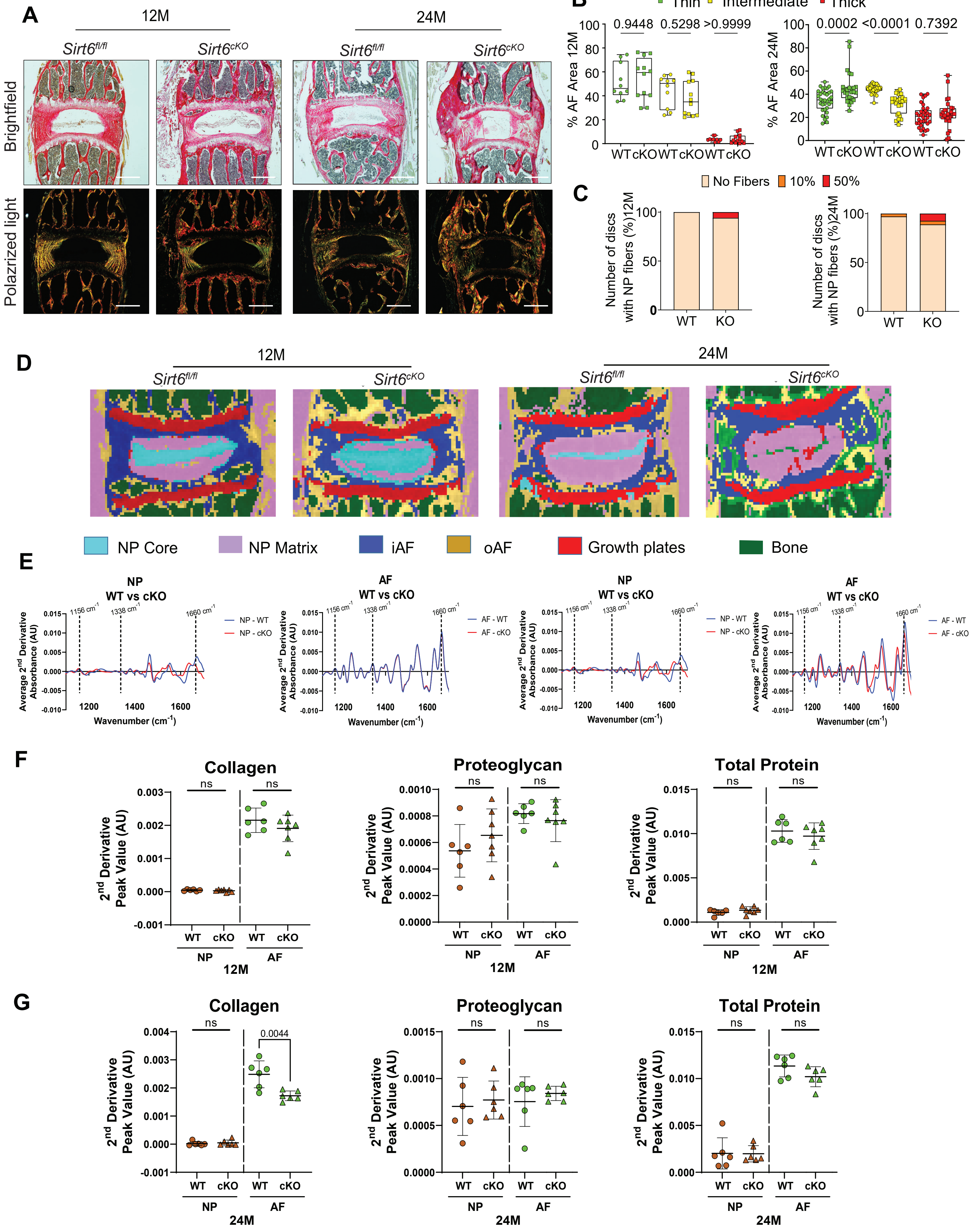

### Supplementary Figure 4

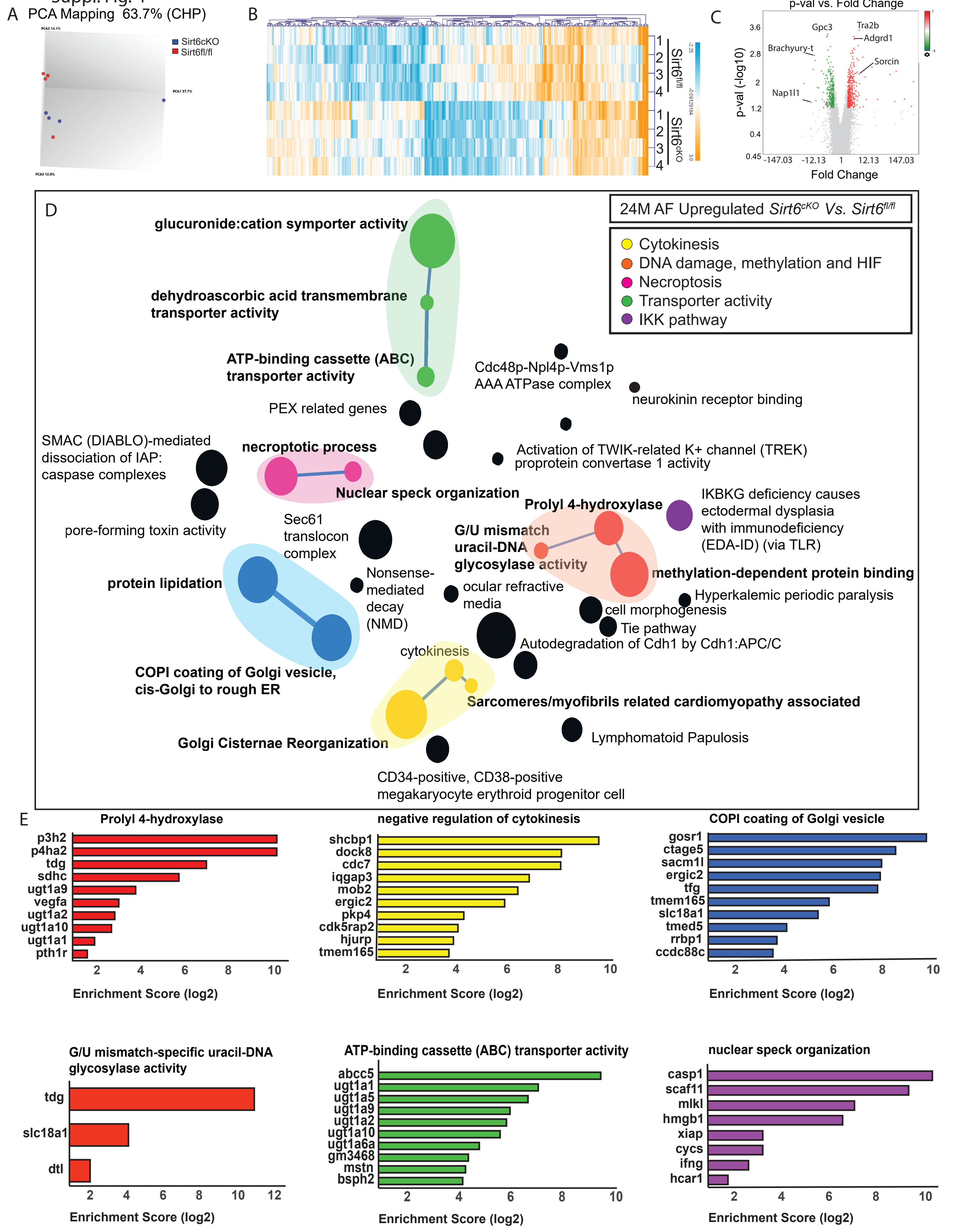

### Supplementary Figure 5

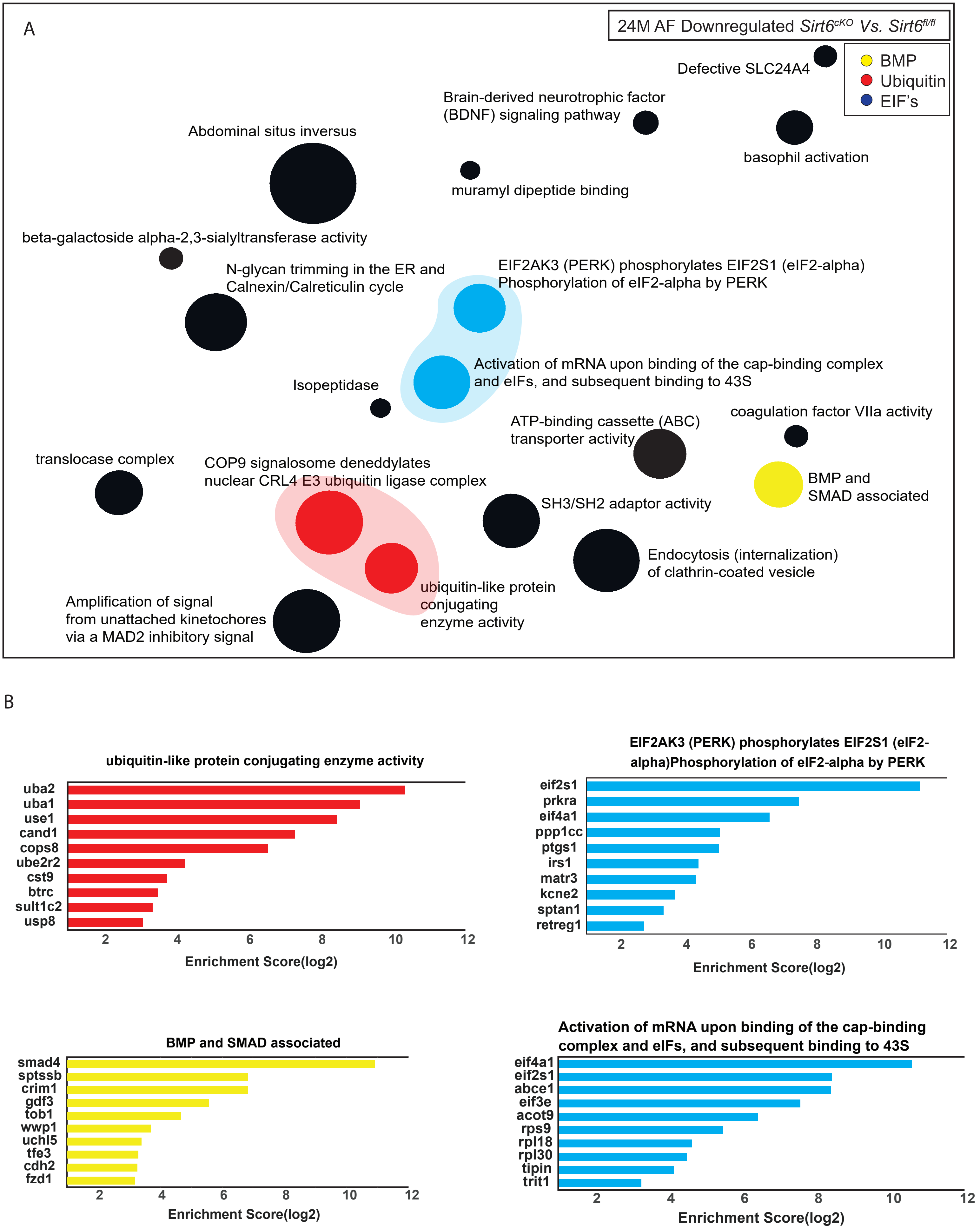

### Supplementary Figure 6

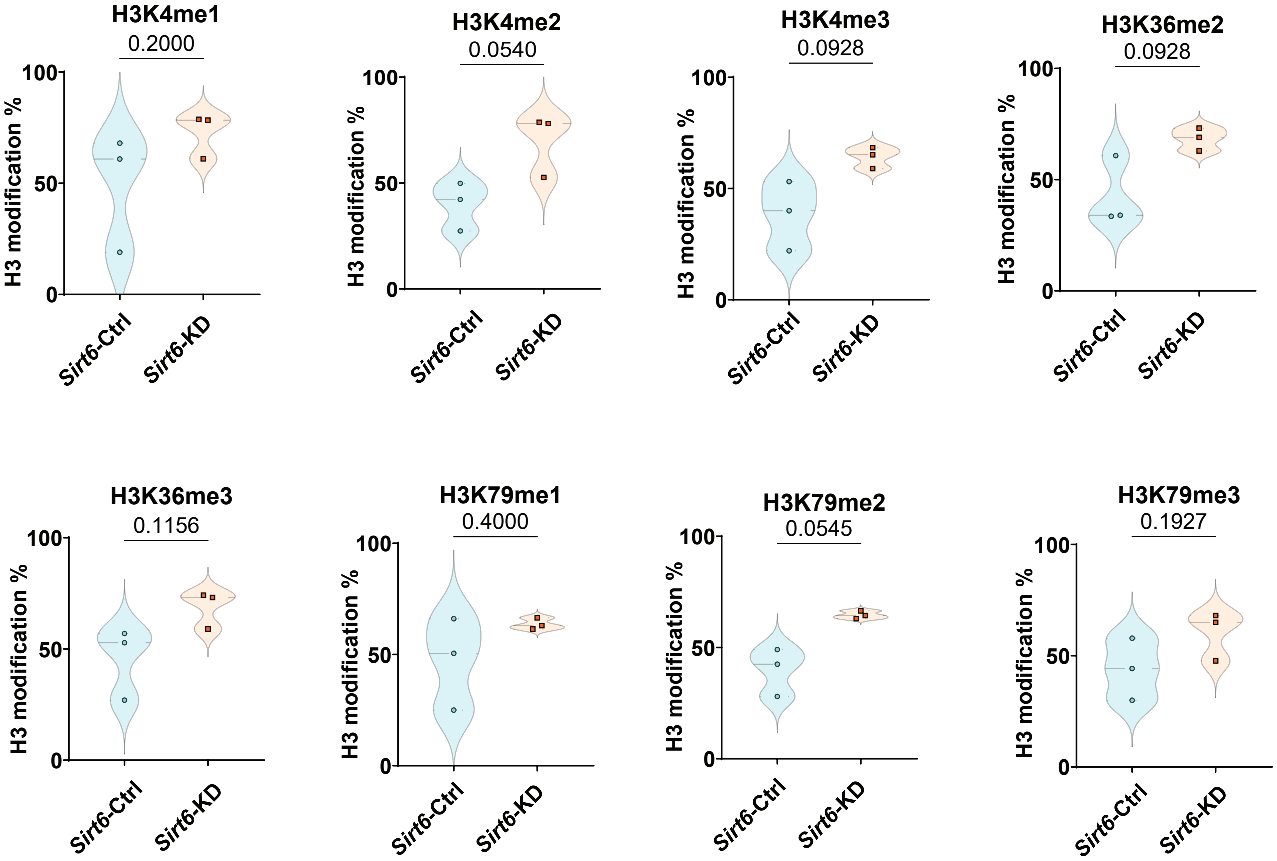

### Supplementary Figure 7

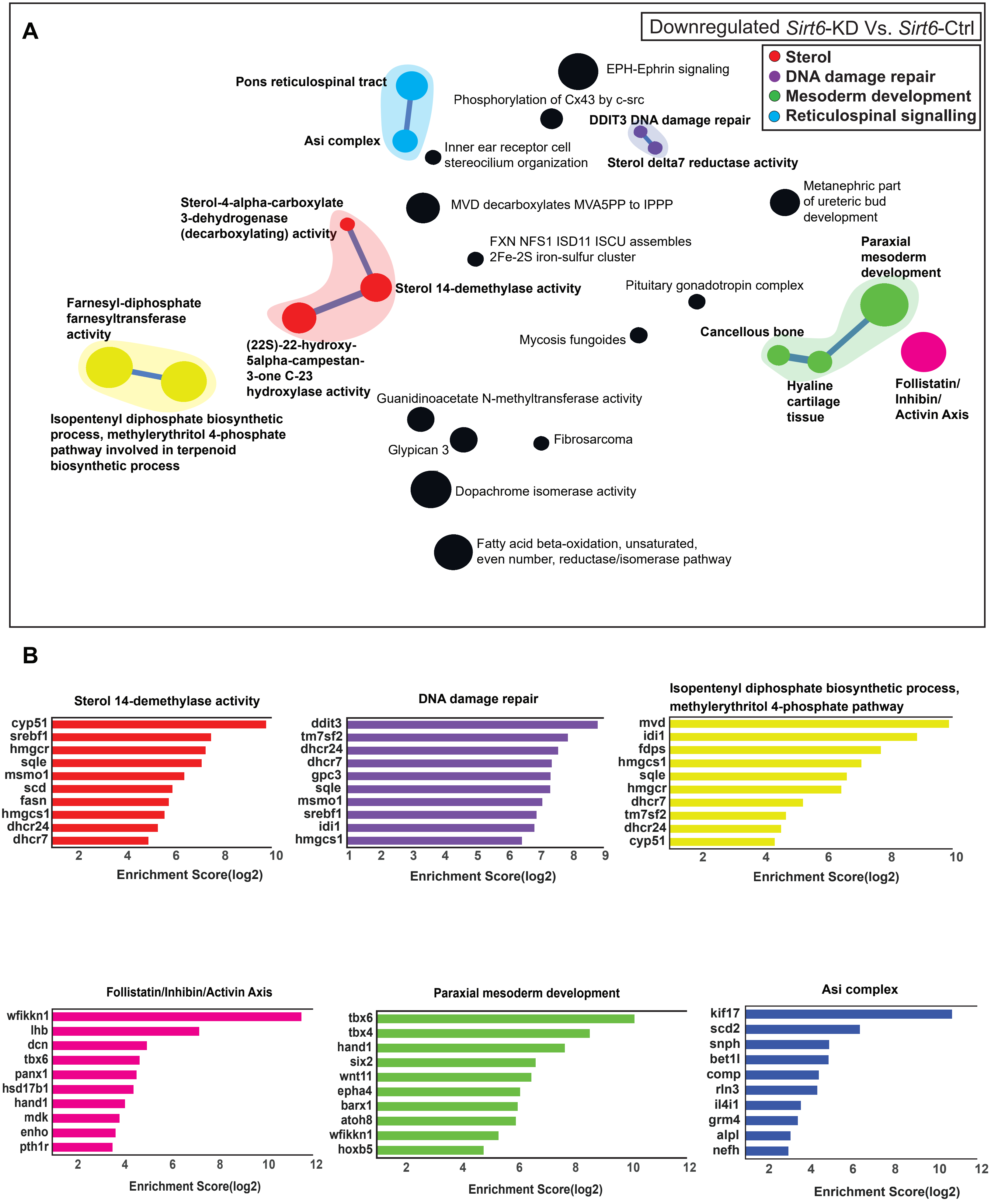

### Supplementary Figure 8

Suppl. Fig. 8

#### Upregulated in *Sirt6*cKO (NP tissue) Vs. Upregulated in *Sirt6*-KD (NP cells)

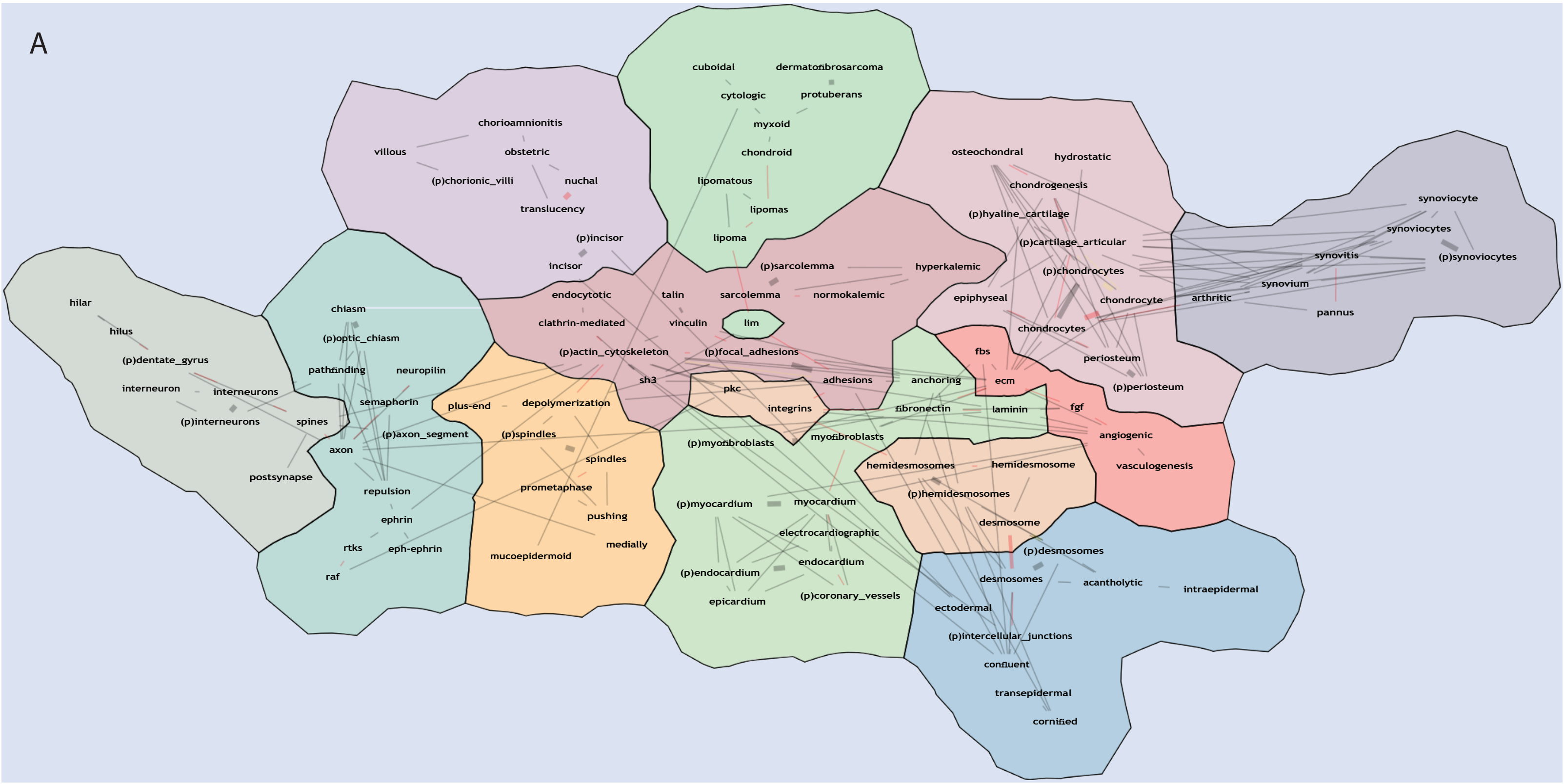

### Supplementary Figure 9

Suppl. Fig. 9

Downregulated in *Sirt6*cKO (NP tissue) Vs. Downregulated in *Sirt6*-KD (NP cells)

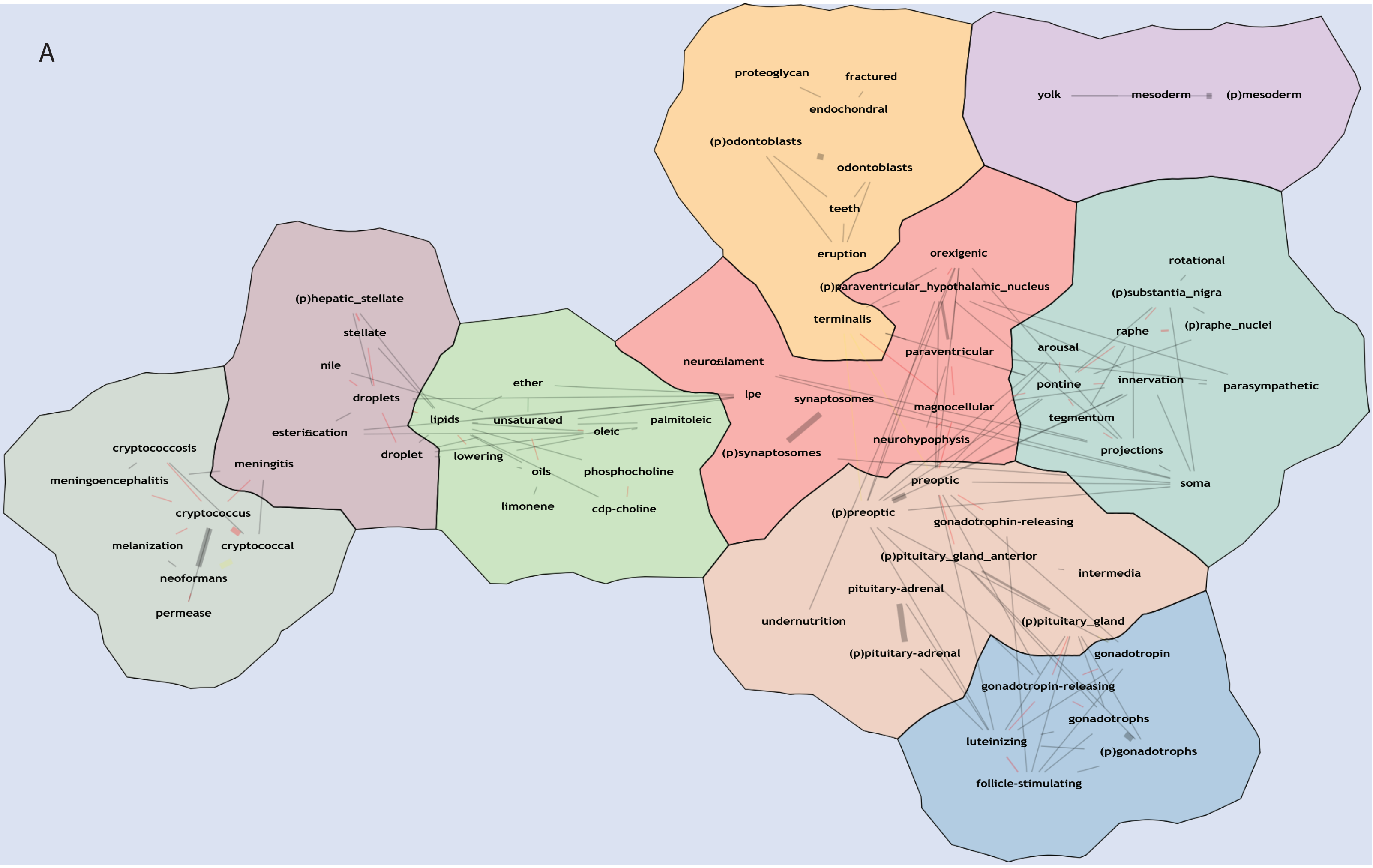

### Supplementary Figure 10

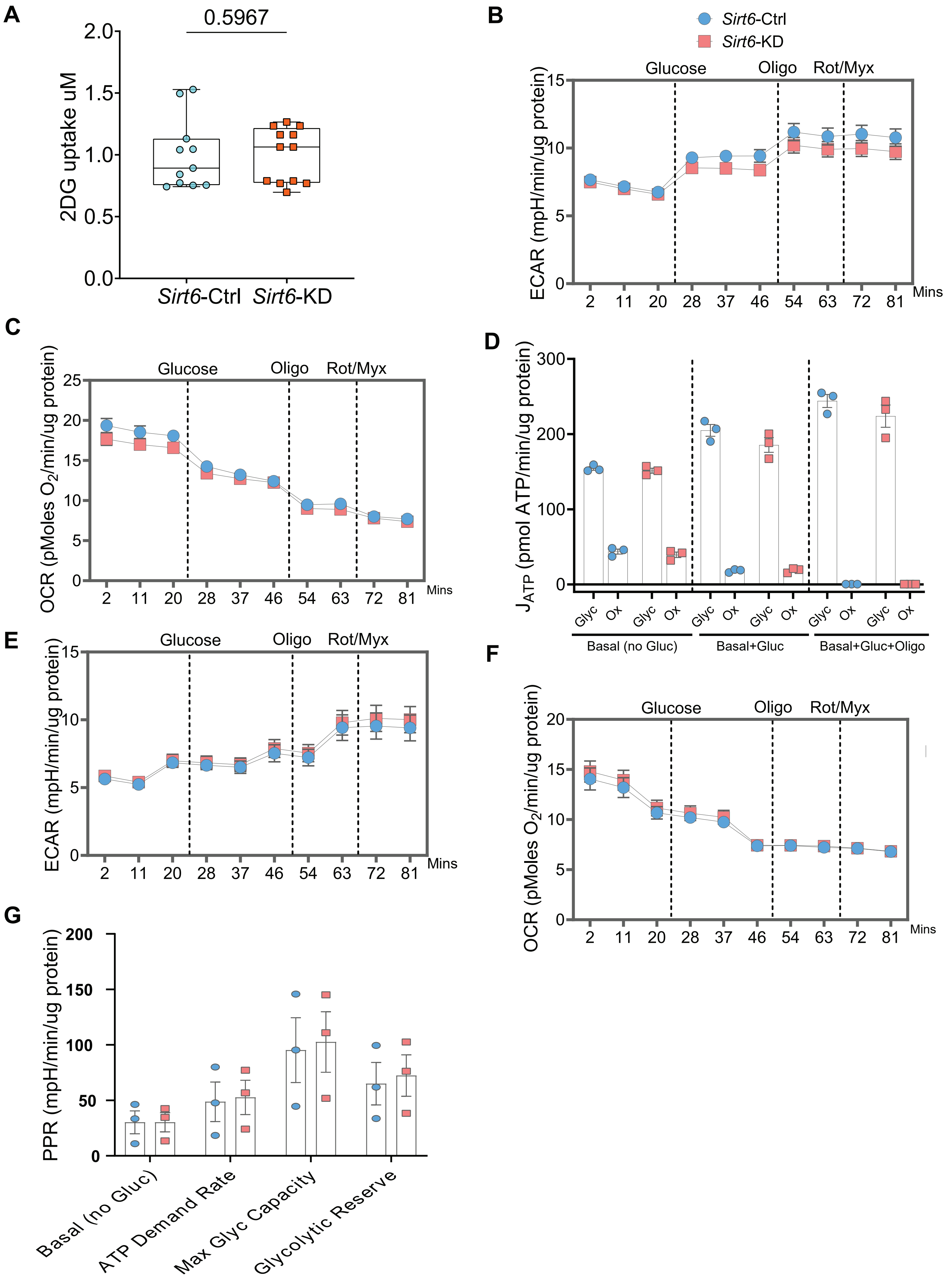
