## Supplementary Figure Legends for "SIRT6 loss causes intervertebral disc degeneration in mice by promoting senescence and SASP status"

**Supplemental Figure Legends:**

**Supplementary Figure 1:** (A) Representative images of Safranin-O/Fast Green staining of *Sirt6*$^{cKO}$ and *Sirt6*$^{fl/fl}$ caudal discs at 12 months and 24 months. Scale bar = 200 μm. (B-C) Distribution and Average Modified Thompson Grades of caudal discs of *Sirt6*$^{cKO}$ and *Sirt6*$^{fl/fl}$ at (B) 12 months and (C) 24 months. N = 3-6 mice/genotype, 3-4 discs/animal. (D-E) Age dependency of Modified Thompson Grades for (D) NP and (E) AF of lumbar discs showing accelerated aging in the AF compartment of *Sirt6*$^{cKO}$ mice. N = 4-6 animals/genotype and 3-4 levels/animal. Statistical difference between grade distributions was tested using chi-square test, all other quantitative data was compared using unpaired t-test, $^*p < 0.05$.

**Supplementary Figure 2:** Representative μCT reconstructions showing (A-B) transverse section through lumbar vertebra and (C-D) lumbar motion segment in 12M and 24M old *Sirt6*$^{fl/fl}$ and *Sirt6*$^{cKO}$ mice. (E) Trabecular bone properties, BV/TV, Tb. Th. (trabecular thickness), Tb. N. (trabecular number), Trab. Sp. (trabecular separation) (F) Cortical bone properties of Cs. Th. (cross-sectional thickness) and closed porosity are shown. Statistical difference between groups was tested using unpaired t-test, $^*p < 0.05$.

**Suppl. Figure 3:** (A) Representative polarized images of Picrosirius Red-stained lumbar disc sections and (B) quantification of collagen fibers from 12 and 24M *Sirt6*$^{fl/fl}$ and *Sirt6*$^{cKO}$ mouse discs. n= 4-6 animals/genotype and 3-4 discs/animal. Scale bar= 200 μM. (C) Quantification of NP compartment fibrosis. (D) Spectral cluster analysis images of 12 M and 24 M discs (scale bar = 200 μm). IVD n=4-6 animals/genotype and 3-4 discs/animal. Scale bar=200uM. (E) Average superimposed second derivative spectra, inverted for positive visualization of the NP and AF of 12M and 24M old mice. (F, G) quantification of mean second derivative peaks for 12M (F) and 24M (G). Significance for quantitative measures was determined by using an unpaired *t*-test with Mann–Whitney test or Welch's correction, as appropriate. Quantitative measurements represent the median with the interquartile range.

**Supplementary Figure 4**: Microarray analysis of AF tissue transcripts from *Sirt6*$^{fl/fl}$ and *Sirt6*$^{cKO}$ represented as (A) Three-dimensional Principal component analysis (PCA) showing discrete clustering of based on genotype (*n* = 4 mice/genotype, 5-6 pooled discs/animal) (B) Heat map and hierarchical clustering of Z-score of differentially expressed genes (DEGs) between *Sirt6*$^{fl/fl}$ and *Sirt6*$^{cKO}$ ($p \leq 0.05$, FC≥1.75). (C) Volcano plot of DEGs in the AF showing *p*-value *versus* magnitude of change (fold change). (D) CompBIO analysis of Upregulated DEGs in AF tissue of 24M *Sirt6*$^{cKO}$ represented in a ball and stick model. The enrichment of themes is shown by the size of the ball and connectedness is shown based on thickness of the lines between them. Themes of interest are colored, and superclusters comprised of related themes are highlighted. (E) Top thematic DEGs plotted based on CompBio entity enrichment score.

**Supplementary Figure 5**: (A) CompBIO analysis of downregulated DEGs in AF tissue of from 24M *Sirt6*$^{cKO}$ mice represented as a ball and stick model (FC≥1.75, p>0.05). n=4 mice/genotype 5-6 pooled discs/animal. The enrichment of themes is shown by the size of the ball and connectedness is shown based on thickness of the lines between them. Themes of interest are colored, and superclusters comprised of related themes are highlighted. (B) Top thematic DEGs plotted based on CompBio entity enrichment score.

**Supplementary Figure 6:** (A) Quantitative ELISA showing Histone 3 modifications in *Sirt6*-KD compared to *Sirt6*-Ctrl NP cells (n=3 independent experiments). Significance for quantitative measures was determined by using an unpaired *t*-test.

**Supplementary Figure 7:** (A) Compbio analysis for downregulated DEGs from RNAseq analysis of *Sirt6*-KD Vs. *Sirt6*-Ctrl NP cells (FC≥1.5, FDR≤0.05), represented as a ball and stick model. The enrichment of themes is shown by the size of the ball and connectedness is shown based on thickness of the lines between them. Themes of interest are colored, and superclusters comprised of related themes are highlighted. (B) Top thematic DEGs plotted based on CompBio entity enrichment score.

**Supplementary Figure 8**: Territorial maps of CompBio Assertion engine analysis showing shared themes between upregulated DEGs from NP tissues of *Sirt6*$^{cKO}$ vs *Sirt6*-KD NP cells, p>0.05.

**Supplementary Figure 9:** Territorial maps of CompBio Assertion engine analysis showing shared themes between downregulated DEGs from NP tissues of *Sirt6*$^{cKO}$ vs *Sirt6*-KD NP cells, p>0.05.

**Supplementary Figure 10**: (A) 2-Deoxyglucose (2-DG) uptake in *Sirt6*-Ctrl Vs. Sirt6-KD NP cells for 24 hours. (B-D) OCR and ECAR traces in *Sirt6*-Ctrl Vs. Sirt6-KD NP cells for 24-hour to assess ATP production rate. (E-G) OCR and ECAR traces in *Sirt6*-Ctrl Vs. *Sirt6*-KD NP cells for 24-hour to assess proton production rate.
